# Striatal spinophilin enhances D2R interaction with cytosolic proteins to mediate persistent D2R agonist-induced locomotor suppression

**DOI:** 10.1101/2025.09.09.675136

**Authors:** Basant Hens, Whitney R. Smith-Kinneman, Emma H. Doud, Anthony J. Baucum

**Author notes:** The Department of Biochemistry and Molecular Biology (Smith-Kinneman, Doud) and Pharmacology and Toxicology (Baucum) are currently undergoing a merger. We have listed the name of the merged department for these individuals.

## Abstract

Loss of dopamine neurons in Parkinson disease (PD) leads to motor deficits. Dopamine D2 receptor (D2R) agonists treat PD-associated motor deficits by acting on postsynaptic receptors located within the striatum that have been upregulated due dopamine loss. However, mechanisms that contribute to increased D2R activity in PD to enhance D2R function are poorly described. Spinophilin is a protein phosphatase 1 targeting protein that is expressed in postsynaptic dendritic spines and interacts with postsynaptic D2Rs. However, how spinophilin regulates D2R function is unknown. In the current study, we found that genetic knockout of spinophilin limited the suppression of locomotion caused by the D2R agonist, quinpirole. Mechanistically, we found that spinophilin is required for quinpirole-induced increases in the interaction of the D2R with intracellular proteins, suggesting spinophilin mediates agonist-induced D2R internalization. Therefore, our data support future studies targeting the spinophilin/D2R interaction to enhance the efficacy of current PD therapeutics.

## INTRODUCTION

Motor function requires highly coordinated and skilled movements. These movements are regulated by a host of interconnected brain regions, such as the striatum. Striatal dysfunction leads to a variety of neurological disorders, including Parkinson disease (PD). The major hallmark of PD is loss of dopamine projections to the striatum. To date, there is no effective cure for PD^1,2^. However, dopamine replacement therapy with levodopa (L-Dopa) or dopamine 2 receptor-family (D2R) agonists are used as pharmacological treatments^3^. Moreover, deep brain stimulation (DBS) has emerged as an effective treatment for some people who are refractory to pharmacological treatments^4^. Unfortunately, dopamine replacement therapies are associated with dyskinesias or/and waning efficacy. Additionally, DBS is highly invasive, not always effective, and can present with severe adverse events^5^. Therefore, there is a critical need to identify striatal targets that may provide insights into novel therapeutic interventions for PD.

Loss of dopamine projections to the striatum will impact downstream function of multiple neurons within the striatum. In the rodent striatum, ∼95% of neurons are medium spiny neurons (MSNs) with the remainder being interneurons. Classically, the two subclasses of striatal MSNs: indirect pathway MSNs (iMSNs) and direct pathway MSNs (dMSNs), inhibit and promote movement, respectively^6^. Molecularly, iMSNs express D2Rs, whereas dMSNs express dopamine D1 receptors (D1Rs)^6^. Importantly, D2Rs inhibit iMSN activity whereas D1Rs enhance dMSN activity. Inhibiting iMSN activity or enhancing dMSN activity promotes movement^6^. In addition to postsynaptic D2Rs, D2Rs are also found presynaptically on glutamate and dopamine terminals within the striatum. In contrast to postsynaptic dopamine receptor activation, presynaptic D2R activation decreases glutamate and dopamine release and limits movement^7^.

In rodent models, the D2R-family agonist quinpirole leads to locomotor suppression that is lost over multiple injections^8^. This loss of locomotor suppression may be due to differential agonist-induced regulation of presynaptic and postsynaptic D2Rs. Specifically, pre- and post-synaptic D2Rs have two unique splicing variants, a long isoform (D2L) and a short isoform (D2S) with the D2L being postsynaptically enriched and the D2S presynaptically enriched^9^. Moreover, the splicing form of the D2R has effects on receptor glycosylation, trafficking, and downstream function, which may reflect differences in how they respond to chronic agonist treatment^10,11^.

One potential candidate that may specifically modulate postsynaptic-enriched D2Rs is the protein phosphatase 1 (PP1)-interacting protein, spinophilin. Spinophilin interacts with postsynaptic D2Rs^12–14^ and has been shown to regulate striatal dopamine-associated behaviors. Specifically, loss of spinophilin enhanced cocaine reward, reduced amphetamine sensitization, and decreased the latency to fall on a rotarod motor learning task^14–16^. Moreover, following dopamine depletion, spinophilin’s binding to PP1 is increased while interactions with myriad other proteins are decreased^17^. Conversely, excessive dopamine activity associated with exposure to psychostimulant drugs of abuse increased spinophilin protein interactions^18^. Together, these data suggest that spinophilin interacts with the D2R and that spinophilin interactions are sensitive to dopamine levels in the striatum. However, the functional regulation of D2Rs by spinophilin is unclear. Using spinophilin global and conditional knockout lines combined with behavioral and biochemical approaches, we found that striatal postsynaptic spinophilin is required for quinpirole-induced increases in D2R intracellular protein interactions and locomotor suppression, suggesting that spinophilin mediates quinpirole-induced changes in D2R trafficking and function. Our results suggest a role for spinophilin in promoting agonist-induced internalization of the D2R and indicate that spinophilin may be a target to enhance efficacy of current PD therapeutics.

## RESULTS

### Spinophilin targets PP1 to D2R in Neuro2A cells

Spinophilin interacts with multiple G-protein coupled receptors, such as opioid receptors, adrenergic receptors, group I metabotropic glutamate receptors, and the D2R^12,19–22^. Of these interactions, the least is known about how spinophilin regulates D2R function. To begin to understand the role of spinophilin in mediating D2R function, we employed a proximity labeling strategy that we have recently developed^23^. Specifically, we created a plasmid encoding the *Drd2* gene that contained a lox-stop sequence followed by a DNA sequence encoding the ALFA epitope-tag^24^ and the UltraID^24^ biotin ligase (**Figure 1A**). To validate appropriate biotin labeling with the construct, we transfected mouse neuro2A cells with this plasmid and either wildtype or an F451A mutant spinophilin, which cannot bind PP1, in the absence or presence of a construct encoding Cre recombinase and probed for streptavidin. We observed a more robust streptavidin signal when Cre-recombinase was co-expressed (**Figure 1B**). Moreover, we observed more robust spinophilin binding in the pulldowns when Cre-recombinase was expressed (arrowheads). We performed additional transfections of mouse neuro2A cells with this plasmid with Cre recombinase in the absence of spinophilin or in the presence of wildtype or an F451A mutant spinophilin. Cells were treated with biotin and a streptavidin pulldown using trypsin-resistant streptavidin was performed. We observed streptavidin staining along with spinophilin expression and pulldown when it was also overexpressed (**Figure S1**). We performed quantitative proteomics using tandem-mass-tag (TMT) labeling. Three samples per group were subjected to TMT labeling and mass spectrometry and abundances for all proteins and peptides detected within each TMT channel are given (**Tables S1 and S2**, respectively). We appended the D2R-ALFA-ultraID (D2R-uID) sequence and the HA-tagged mutant human spinophilin sequence to the mouse protein database search. Across the samples, we detected 179 peptide spectral matches (PSMs) against D2R-uID and 38 PSMs against spinophilin (**Table S1**). The abundance for D2R-uID was fairly consistent across groups (**Figure 1C)**. Moreover, spinophilin expression was greater when either wildtype (WT) or F451A mutant spinophilin was co-expressed (**Figure 1D**). Next, we normalized the expression of spinophilin to total D2R-uID within the same channel and found a similar pattern (**Figure 1E**). In addition to spinophilin, we evaluated the expression of the 3 PP1 isoforms within each group and qualitatively observed greater overall expression of PP1 when WT, but not F451A mutant spinophilin was overexpressed (**Figure 1F**).

**Figure 1.**
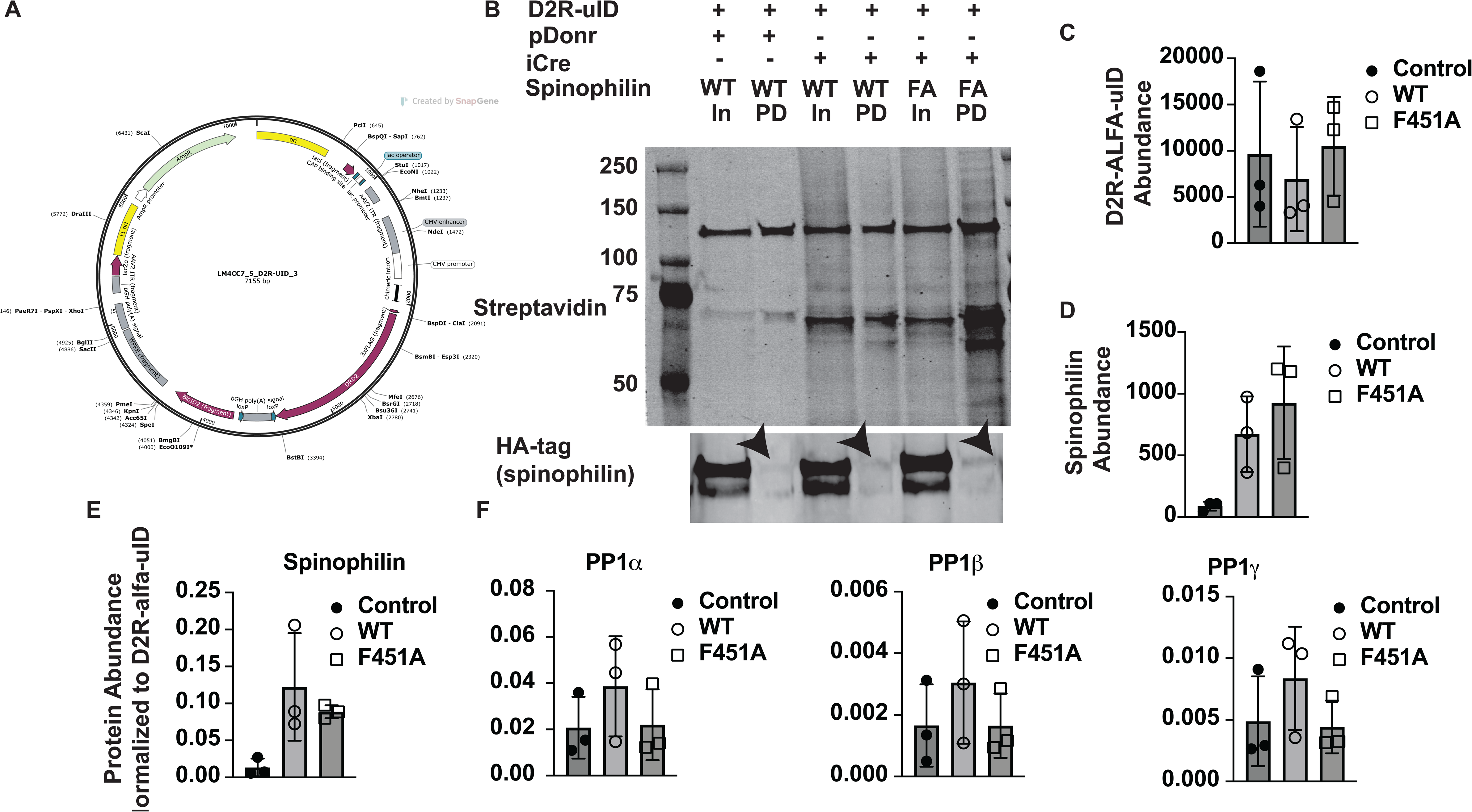
Proteomic identification of spinophilin and PP1 in D2R-UltraID-expressing neuro2A cells. **A.** Sequence validation of the plasmid encoding the long-isoform, human D2R cDNA, a 3’ floxed stop site, an ALFA tag and the ultraID (uID) sequence (D2R-uID). **B.** Lysate inputs (In) or neutravidin pulldowns (PD) from Neuro2A cells expressing D2R-uID and pDonr empty plasmid, cre-recombinase, and/or HA-tagged wildtype (WT) or F451A mutant spinophilin (FA) were blotted with streptavidin or an HA antibody. **C-F**. Trypsin resistant streptavidin pulldowns from Neuro2A cells expressing D2R-uID alone or D2R-uID along with HA-tagged wildtype (WT) or F451A mutant spinophilin (FA) were subjected to tandem mass tag (TMT) proteomics. Total abundance matching D2R-uID (**C**) and HA-tagged spinophilin (**D**) are given. Abundance values matching HA-tagged spinophilin (**E**) and PP1 isoforms (**F**) were further normalized to the abundance of the D2R-uID within the corresponding sample. N=3 independent transfections per condition.

### Identification of the D2R interactome

The proteomics analyses identified 1,234 total proteins across all channels (**Tables S3A**). Of those, 956 were not contaminants or known endogenously biotinylated proteins (e.g. carboxylases) and had 4 or more total PSMs detected (**Table S3B**). Of these, 945 proteins were mapped by STRING database (STRING-db) for protein-protein and network analyses^25–28^. The top 10 gene ontology (GO) pathways for cellular component (**Figure S2A**), molecular function (**Figure S2B**), and biological process (**Figure S2C**) revealed that the interactome matched multiple ribosomal, translation, mitochondrial, and metabolic proteins. Moreover, consistent with the D2R being a target for current PD therapeutics, the PD pathway was the second most enriched pathway in the in the Kyoto Encyclopedia for Genes and Genomes (KEGG) analysis (**Figure S2D**). A full list of significant pathways detected are given in **Table S4**.

### Spinophilin enhances the D2R interactome in neuro2A cells

Our proximity labeling data suggest that the D2R interacts with myriad proteins in a neuronal cell line, including proteins associated with PD. A caveat to these results is that proximity labeling studies may lack selectivity in that overexpression of any protein fused to UltraID will interact with proteins involved in protein homeostasis as well as high abundance proteins. Therefore, this may be the case with the ribosomal proteins, mitochondrial proteins, and proteasomal proteins that we detected. To enhance functional relevance, to delineate D2R interacting proteins that may be impacted by spinophilin, and to identify how spinophilin regulates D2R function in vivo, we evaluated how spinophilin overexpression impacts the D2R interactome. Interestingly, when evaluating total protein expression upon WT spinophilin overexpression, we found that there was an overall greater abundance of proteins associated with the D2R (**Figure 2A**). We visualized the top 50 interacting proteins by log_2_-fold change in WT overexpressing cells compared with non-spinophilin expressing cells using the STRING-db and found that a majority of them were mitochondrial proteins (**Figure 2B**). Pathway analysis of this extended list included mitochondrial proteins and functions (**Figure 2C-F**). Moreover, as with the total interactome, the KEGG pathway analysis also included PD (**Figure 2F**). The full complement of pathway enrichments is given in **Table S5**. Therefore, spinophilin overexpression seems to stabilize D2R protein interactions and has a robust effect on the interaction of D2R with mitochondrial and other intracellular proteins and processes.

**Figure 2.**
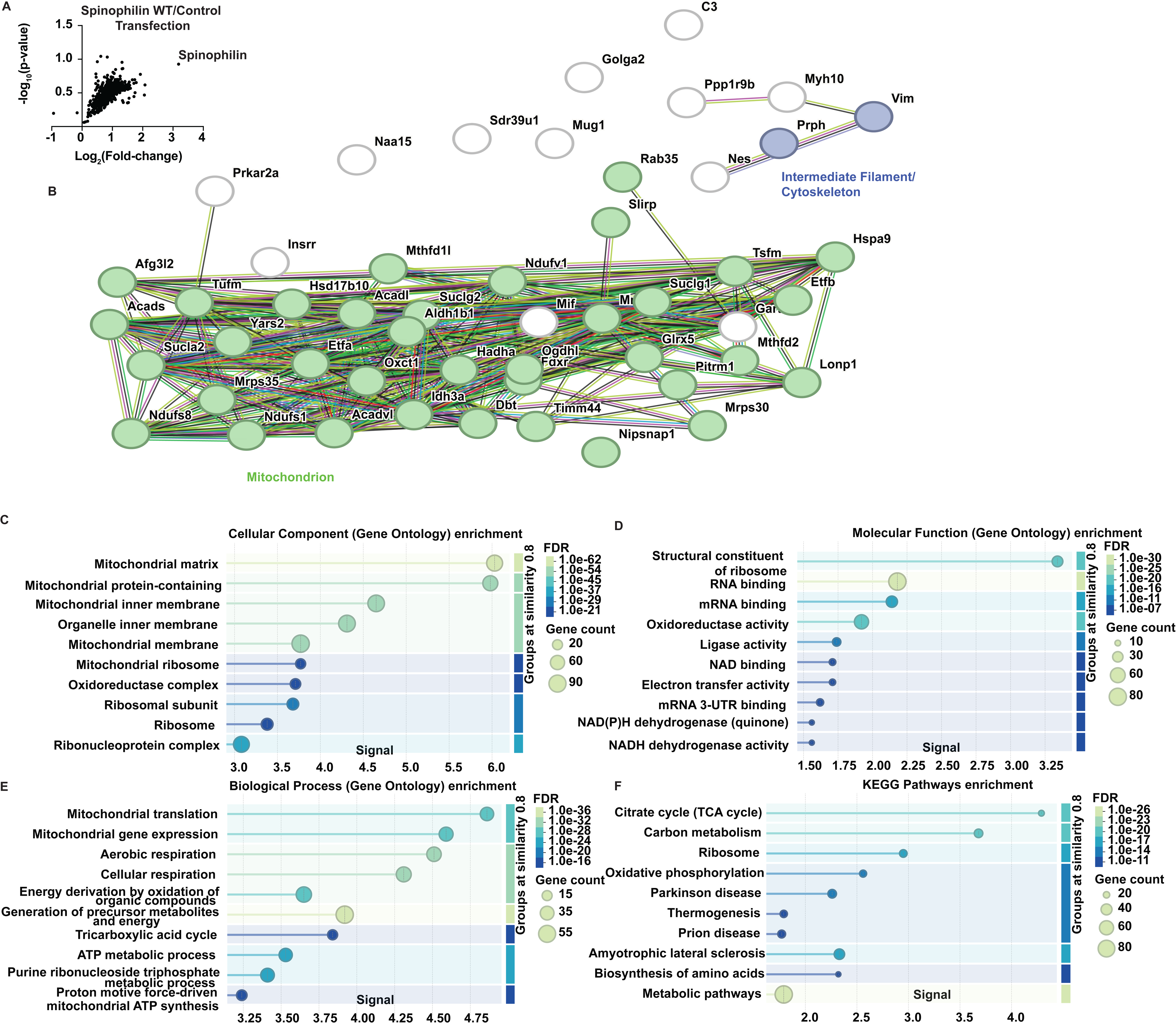
Spinophilin overexpression enhances D2R interactions with intracellular proteins in neuro2A cells. **A.** Volcano plot showing increased protein abundance in streptavidin pulldowns from cells expressing D2R-uID and WT spinophilin compared with D2R-uID expressing cells alone. **B.** String-db map of the top 50 most enriched interacting proteins by fold-change in streptavidin pulldowns from cells expressing D2R-uID and WT spinophilin compared with D2R-uID expressing cells alone. **C-F**. Proteins with log_2_-fold change > 1 were input into STRING-db and the top 10 gene ontology (GO) **C)** cellular component, **D)** molecular function, and **E)** biological process, and **F)** Kyoto Encyclopedia for Genes and Genomes (KEGG) pathways are listed. N=3 independent transfections per condition.

### Spinophilin association with PP1 is required for enhancing D2R interactions in neuro2A cells

As stated above, many of the proteins that the D2R is interacting with may be non-selective. Moreover, expression of any additional protein may impact cell health and perturb any protein interactome. Therefore, we compared the D2R interactome in the presence of overexpressed mutant spinophilin (F451A), that does not bind PP1, and the non-spinophilin expressed condition. Unlike WT spinophilin, F451A mutant spinophilin had limited effects on altering the D2R interactome (**Figure 3A**). Therefore, PP1-binding is required for the spinophilin-dependent increase in D2R protein interactions. This is further demonstrated when comparing the D2R interactome in the presence of F451A mutant spinophilin compared with WT spinophilin as overall there is decreased abundance of proteins in the mutant compared with WT conditions (**Figure 3B**). When comparing the top 50 most decreased proteins in the mutant compared with WT spinophilin overexpressing cells, we observed multiple mitochondrial, intermediate filament, and vesicle trafficking/Golgi proteins (**Figure 3C**). Moreover, when evaluating proteins with a log_2_-fold change of -1 or lower in the mutant, compared with WT, spinophilin, we found that similar classes of proteins were decreased as were increased in the WT overexpression compared to no spinophilin overexpression conditions. These include proteins associated with the mitochondrion and multiple mitochondrial processes and ribosomal proteins. The top 10 GO enrichments within the cellular component, molecular function, and biological process are shown (**Figure 3D-F**). We also observed PD within the top 10 KEGG pathways (**Figure 3G**). Full enrichments are given in **Table S6.**

**Figure 3.**
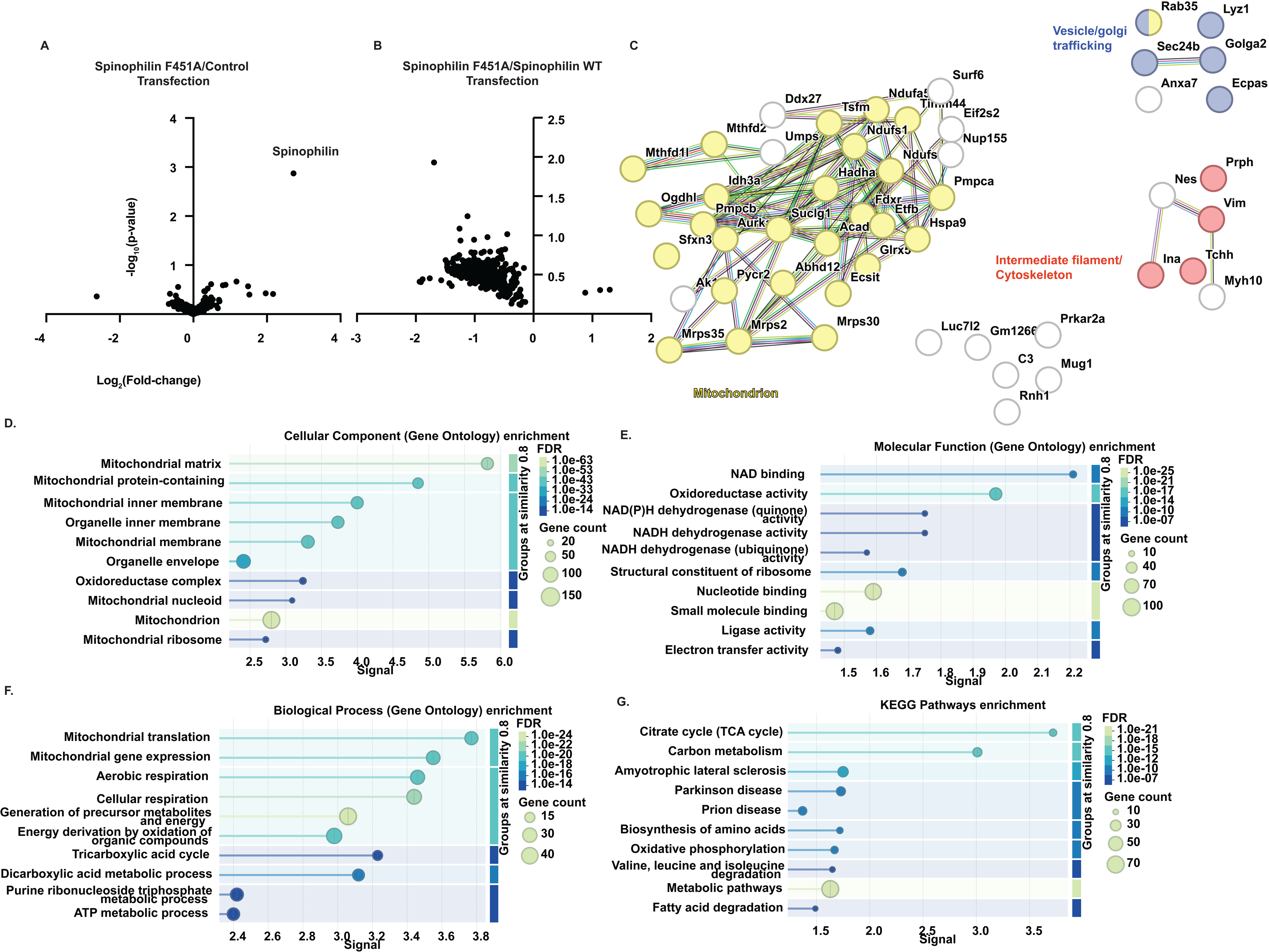
Spinophilin’s association with PP1 is required for enhancing D2R interactions within intracellular proteins in neuro2A cells. **A**. Volcano plot showing minimal difference between F451A mutant spinophilin and minimally altered the D2R interactome. **B**. Volcano plot showing decreased protein expression in the streptavidin pulldowns from mutant F451A spinophilin-expressing cells compared with WT spinophilin expressing cells. **C**. String-db map of the top 50 most decreased proteins in streptavidin pulldowns isolated from F451A mutant spinophilin compared with WT spinophilin overexpressing cells. **D-F**. Proteins with log_2_-fold change < 1 were input into STRING-db and the top 10 gene ontology (GO) for **D)** cellular component, **E)** molecular function, and **F)** biological process, and the **G)** Kyoto Encyclopedia for Genes and Genomes (KEGG) are given. N=3 independent transfections per condition.

### Spinophilin mediates quinpirole-induced hypolocomotion

Chronic treatment with the D2R agonist quinpirole has complex actions on motor output. For instance, chronic administration of 3 mg/kg quinpirole to rats daily for 8 days decreased locomotion for the first 3 days and increased locomotion above vehicle injections after day 5^29^. To determine how different doses impact quinpirole-induced locomotion in mice, we injected mice daily for 5 days with saline vehicle or 1, 3, or 10 mg/kg quinpirole. Mice were placed in an open field (Phenotyper cages) and distance traveled was measured using video recordings for one hour. A 2-way repeated measure ANOVA revealed significant time (p=.01) and treatment (p=.0012) effects. When we ran Dunnett’s multiple comparison test, the 3 mg/kg dosage maintained the most consistent decrease in locomotion across the 5 days and showed the largest effect size in terms of hypo-locomotion (day 1: p=.0335, day 2: p=.0086, day3 : p=.0304, day 4: p=.0007, day 5: p=.07) (**Figure S3**). Therefore, we evaluated how loss of spinophilin impacted locomotor suppression to a 3 mg/kg dose of quinpirole.

The D2R-family agonist, quinpirole, causes locomotor suppression, whereas loss of spinophilin, basally, enhances motor output^30^. However, loss of spinophilin attenuated amphetamine-induced locomotor sensitization and excessive motor output in mouse models of repetitive grooming^30,31^. Therefore, spinophilin may have unique effects on locomotion depending on perturbations in dopamine signaling. To determine if spinophilin impacts quinpirole-induced motor suppression, we pre-injected WT (Spino^+/+^) or spinophilin knockout (Spino^-/-^) mice with saline for two days followed by five daily injections of saline vehicle or 3 mg/kg of quinpirole (**Figures 4A and S4**). A 3-way ANOVA found day (p<0.0001), genotype (p=0.0025), day x treatment (p=<0.0001), day x genotype (p=0.0448), and day x treatment x genotype (p=.0071) effects (**Figure 4B**). To further identify the specific components of the interaction effect, we analyzed the data using a 2-way ANOVA analysis within the treatment and genotype groups. When comparing the treatment of saline and quinpirole in Spino^+/+^ animals, we found significant treatment (p=0.0063) and day effects (p=0.001). When comparing the saline and quinpirole groups within Spino^-/-^ mice, we found day (p=0.0004) and day x treatment interaction (p=0.0006) effects. Additionally, we examined the effect of genotype within the quinpirole treatment group and observed day (p<0.0001), genotype (p<0.0082), and day x genotype interaction (p=0.0057) effects.

**Figure 4.**
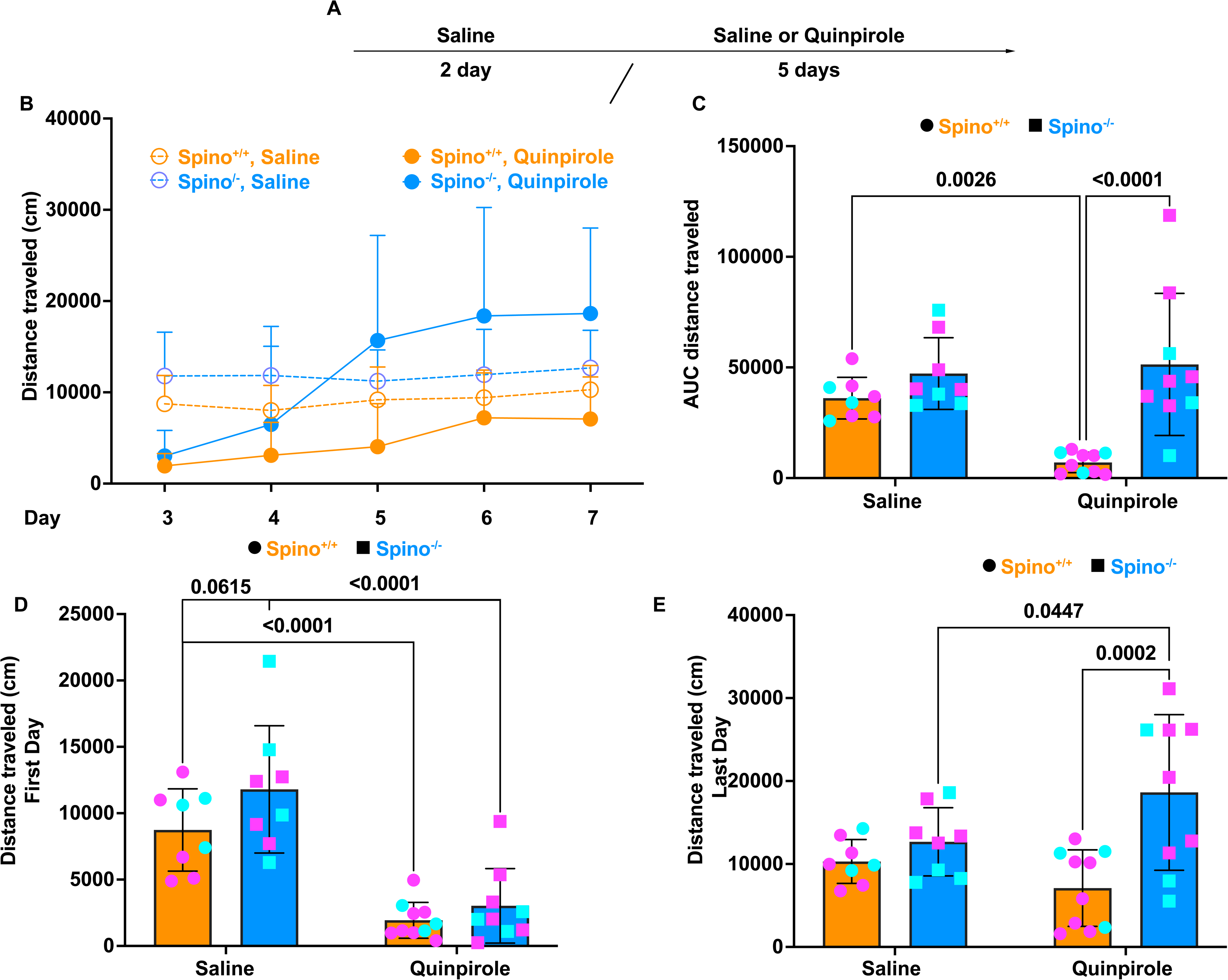
Loss of spinophilin globally abrogated the locomotor suppression of quinpirole. **A.** Spino^+/+^ and Spino^-/-^, 2–8-month-old male or female mice were intraperitoneally injected with saline, daily for 2 days. Subsequently, mice were injected with saline or quinpirole daily for 5 days. The total distance traveled was recorded over 1 hour using the open-field test. **B.** Distance traveled (cm) measurements across injection days 3-7 are plotted. A 3-way ANOVA revealed significant day (p=0.0001), genotype (p<0.0025), day x treatment (p=.0001), and treatment x genotype x day (p=0.0071) effects. **C.** Area under the curve (AUC) analysis for each individual animal was calculated and a 2-way ANOVA was performed. There were significant genotype (p<0.0001) and treatment x genotype (p=0.0139) effects. An uncorrected Fisher’s least significant difference (LSD) post-hoc test showed a significantly greater AUC of distance traveled in Spino^-/-^ mice treated with quinpirole compared to Spino^+/+^ animals (p<0.0001). Moreover, there was significant decrease in the AUC of distance traveled in Spino^+/+^ mice treated with quinpirole compared to saline (p=.0026). **D.** A 2-way ANOVA on the distance traveled on the first day was performed. There was a significant treatment effect (p<.0001). An uncorrected Fisher’s LSD post-hoc test demonstrated a significant suppression of locomotion in quinpirole treated Spino ^+/+^ and Spino^-/-^ mice compared with saline treated animals (p<. 0001, p<.0001). **E.** A 2-way ANOVA on the distance traveled on the last day was performed. There were significant genotype (p=.0014) and genotype x treatment interaction (p=0.0280) effects. An uncorrected Fisher’s LSD post-hoc test showed an increased locomotion in the quinpirole-treated Spino^-/-^ mice compared with the Spino^+/+^ mice (p=.0002). Moreover, there was a greater locomotion in the quinpirole-treated Spino^-/-^ mice compared to saline treated Spino^-/-^ animals (p=0.0447). Mean + (B) or +/-standard deviation is shown. N=8-10 per group. Magenta colors represent female mice, cyan colors represent male mice.

To further quantify the effects, we measured the area under the curve (AUC) for each animal followed by a 2-way ANOVA analysis of the AUCs (**Figure 4C**). We detected genotype (p=0.0001) and treatment x genotype interaction (p=0.0139) effects. An uncorrected Fisher’s least significant difference (LSD) post hoc analysis of the AUC found a significant increase in movement in global Spino^-/-^ mice compared to Spino^+/+^ mice treated with quinpirole (p<0.0001). Additionally, the post hoc analysis demonstrated a significant decrease in movement of Spino^+/+^, but not Spino^-/-^, mice treated with quinpirole compared to saline (p=0.0026).

We next compared how spinophilin impacts locomotor changes following a single or 5-daily injections of quinpirole. We analyzed the total distance traveled on either the first or last day of injections using a 2-way ANOVA followed by an uncorrected Fisher’s LSD analysis (**Figure 4D**). For the first day, there was a significant treatment effect (p<.0001) and a trending genotype effect (p=0.0616). On the first day of treatments, there was a significant reduction in distance traveled between the quinpirole compared to saline treatment in both Spino^+/+^ and Spino^-/-^ animals (p<0.0001). Conversely, a 2-way ANOVA performed on distance traveled on the last day revealed no treatment effect (p= 0.4966), but genotype (p=0.0014) and genotype x treatment interaction (p=0.0280) effects (**Figure 4E**). For the multiple comparisons on the last day in the total distance traveled among groups, there was a significant increase in distance traveled in the Spino^-/-^ compared to Spino^+/+^ mice treated with quinpirole (p=0.0002). Additionally, there was a significant *increase* in the distance traveled in quinpirole treated Spino^-/-^ mice compared to saline-treated Spino^-/-^ mice when performing a within-genotype comparison (p=0.0447). From these findings, our data suggest that loss of spinophilin globally led to hyperlocomotion in quinpirole treated animals at later injection days, suggesting spinophilin is mediating the persistent locomotor suppressive effects of quinpirole.

### Spinophilin regulates quinpirole-induced changes in D2R protein interactions

While spinophilin is mediating D2R agonist-induced locomotor suppression, how it is doing this is unclear. To identify putative mechanisms of spinophilin-dependent regulation of D2Rs, we performed an unbiased, quantitative proteomics analysis of D2R co-immunoprecipitating proteins from Spino^+/+^ and Spino^-/-^ mice treated with saline or quinpirole. One hour following the last saline or quinpirole injection, striata were dissected and lysates were immunoprecipitated for D2R using an antibody to the long isoform of the D2R. Immunoprecipitates were digested with trypsin and underwent TMT labeling and mass spectrometry. In total, we detected 1,367 total proteins across all samples (**Table S7**). There were 1,005 total proteins when removing contaminants, including keratin and immunoglobulins, and proteins with fewer than 4 PSMs (**Table S8**). Of the 1,005 proteins, 994 mapped in the STRING-db. When these proteins were analyzed by pathway analysis, similar to our neuro2A cell data, we observed proteins involved in mitochondrial processes, ribosomes, and translation (**Figure S5A-C**). Moreover, one of the major KEGG pathways that matched to the interactome was PD (**Figure S5D**). When plotting the top 50 most abundant proteins by PSMs, we observed multiple proteins associated with cytoskeleton, mitochondrion, and vesicle-mediated transport (**Figure S5E**). The full enrichment list is given in **Table S9**. Therefore, there is overlap in the protein interactions detected in our neuro2A cell proximity labeling studies and in our D2R immunoprecipitation studies.

Previous studies have found that treatment with quinpirole leads to internalization of the D2R^32–34^. Therefore, we wanted to delineate the effects of quinpirole treatment on the D2R interactome. When evaluating expression of the D2R, we observed fairly equal abundance across all groups, suggesting an even immunoprecipitation of the receptor (**Figure S6A**). We normalized proteins to the expression of the D2R (**Table S10**) and quantified the log_2_-fold change of the interactome across different groups. We found that, overall, quinpirole enhanced the association of proteins that co-immunoprecipitated with the D2R (**Figure S6B**). When evaluating the 847 proteins that had a log_2_-fold increase of 0.5 or greater in the quinpirole-treated, compared with saline-treated, wildtype mice and were present in the STRING-db, we observed GO and KEGG pathway enrichment of proteins associated with the cytoplasm, cytoskeleton binding, and PD (**Figure S6C-F**). When evaluating the top 50 proteins based on increased log_2_-fold change in the quinpirole-treated group, we observed multiple cytoplasmic, cytoskeletal, and mitochondrial proteins (**Figure S6G**). The full enrichment list is given in **Table S11**. This is consistent with an increased interaction with cytoplasmic proteins in the quinpirole-treated samples, further suggesting that quinpirole treatment leads to D2R internalization^33,35^.

Given the difference in locomotor output between Spino^+/+^ and Spino^-/-^ mice within the quinpirole treated group, we decided to compare the abundance of D2R interacting proteins between these two groups (**Table S10**). Consistent with a role for overexpression of spinophilin in Neuro2A cells *enhancing* D2R interactions, we observed an overall *decrease* in the interactions in the Spino^-/-^ compared to Spino^+/+^ quinpirole-treated mice (**Figure 5A**). While variable, there was a qualitatively lower level of PP1 isoforms in the Spino^-/-^ samples, potentially suggesting decreased targeting of PP1 to the D2R (**Figure 5B**). We next input proteins that had an overall decrease of -0.5 or lower in the log_2_-fold change in Spino^-/-^ compared to Spino^+/+^ quinpirole-treated mice into the STRING-db. There were 261 proteins with this criterion that matched the STRING-db. We evaluated the top 10 GO and KEGG enrichments and found overlapping enrichments with our neuro2A cell data, including mitochondrial process, cytoskeletal, vesicle trafficking, and PD pathway proteins (**Figure 5C-F**). The full enrichment list is given in **Table S12**. We also identified terms associated with the synapse, which is consistent with structural differences in a neuron compared to an immortalized, undifferentiated cell line. When evaluating the top 50 interacting proteins, we observed multiple mitochondrial, synaptic, and PD-associated proteins (**Figure 5G**).

**Figure 5.**
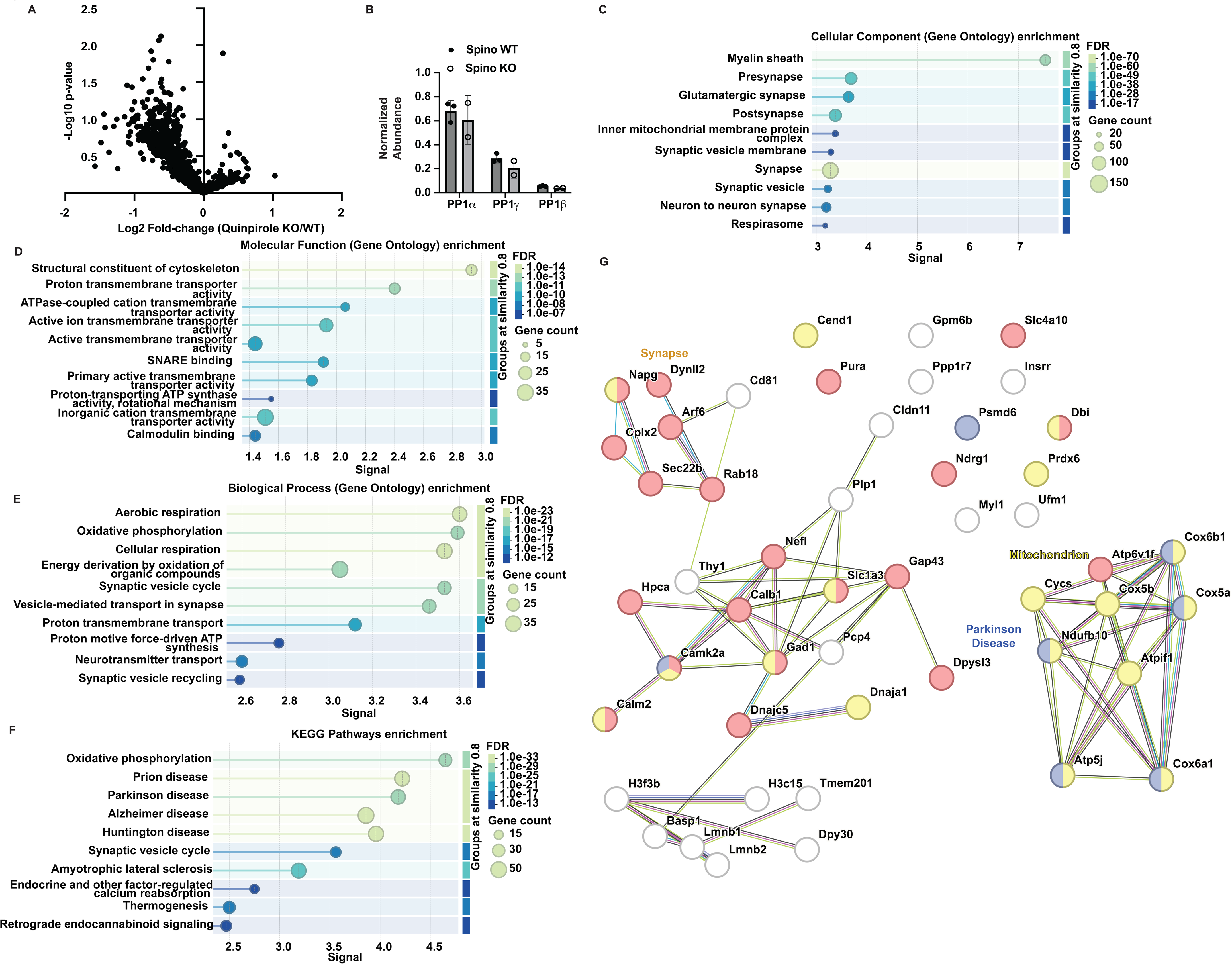
Loss of spinophilin decreased D2R interactions with intracellular proteins. The long-isoform of the D2R was immunoprecipitated from Spino^+/+^ or Spino^-/-^ quinpirole-treated mice and immunoprecipitates were subjected to tandem mass tag proteomics analysis. **A.** Volcano plot showing decreased protein expression in the D2R immunoprecipitations from quinpirole-treated Spino^-/-^ compared with quinpirole-treated Spino^+/+^ mice. **B.** Abundance of PP1 isoforms in the D2R immunoprecipitates in Spino+/+ and Spino^-/-^ mice. **C-F**. Proteins with a log_2_-fold change of -0.5 or lower were input into the STRING-db and a pathway analysis was performed. The top 10 gene ontology pathways for **C)** for cellular component, **D)** molecular function, and **E)** biological process, and the top 10 **F)** Kyoto Encyclopedia for Genes and Genomes (KEGG) pathways are shown. **G)** String-db map of the Top 50 proteins with the greatest decreased log_2_-fold change in Spino^-/-^ compared with Spino^+/+^ mice are given. N=2 Spino^-/-^ and N=3 Spino^+/+^. Mean +/-standard deviation is shown.

### Spinophilin in iMSNs contributes to quinpirole-induced locomotor suppression

Global knockout of spinophilin limited quinpirole-induced locomotor suppression. Moreover, our striatal proteomics suggest that quinpirole-dependent increases in D2R protein interactions with cytoplasmic proteins require spinophilin. Therefore, we wanted to determine how loss of spinophilin from specific striatal neurons impacts quinpirole-induced locomotor changes. To this end, we evaluated locomotor response in our recently described mice that had spinophilin conditionally knocked out of iMSNs (Spino^ΔiMSN^) or littermate controls (Spino^fl/fl^). Mice were injected with saline daily for 2 days and then either saline or quinpirole daily for 5 days. Distance traveled was recorded (**Figures 6A and S7**) and analyzed with a 3-way ANOVA statistical test. We observed significant day (p=0.0004), treatment (p<0.0001), day x treatment (p=.0004), and treatment x genotype (p=0.0257) effects. To further identify the specific components of the interaction effects, we performed a 2-way ANOVA analysis among different groups. When comparing the treatment of saline and quinpirole within Spino^fl/fl^ animals, we found a significant treatment effect (p<0.0001) and day x treatment (p=0.0301) effect. When comparing the treatment of saline and quinpirole in the Spino^ΔiMSN^ mice, we found treatment (p=0.0070), day (p=0.0036), and day x treatment (p=.0195) effects. Additionally, we examined the treatment of quinpirole on the two genotypes, Spino^fl/fl^ and Spino^ΔiMSN^ mice. We found day (p<0.0001) and genotype (p=0.0016) effects.

**Figure 6.**
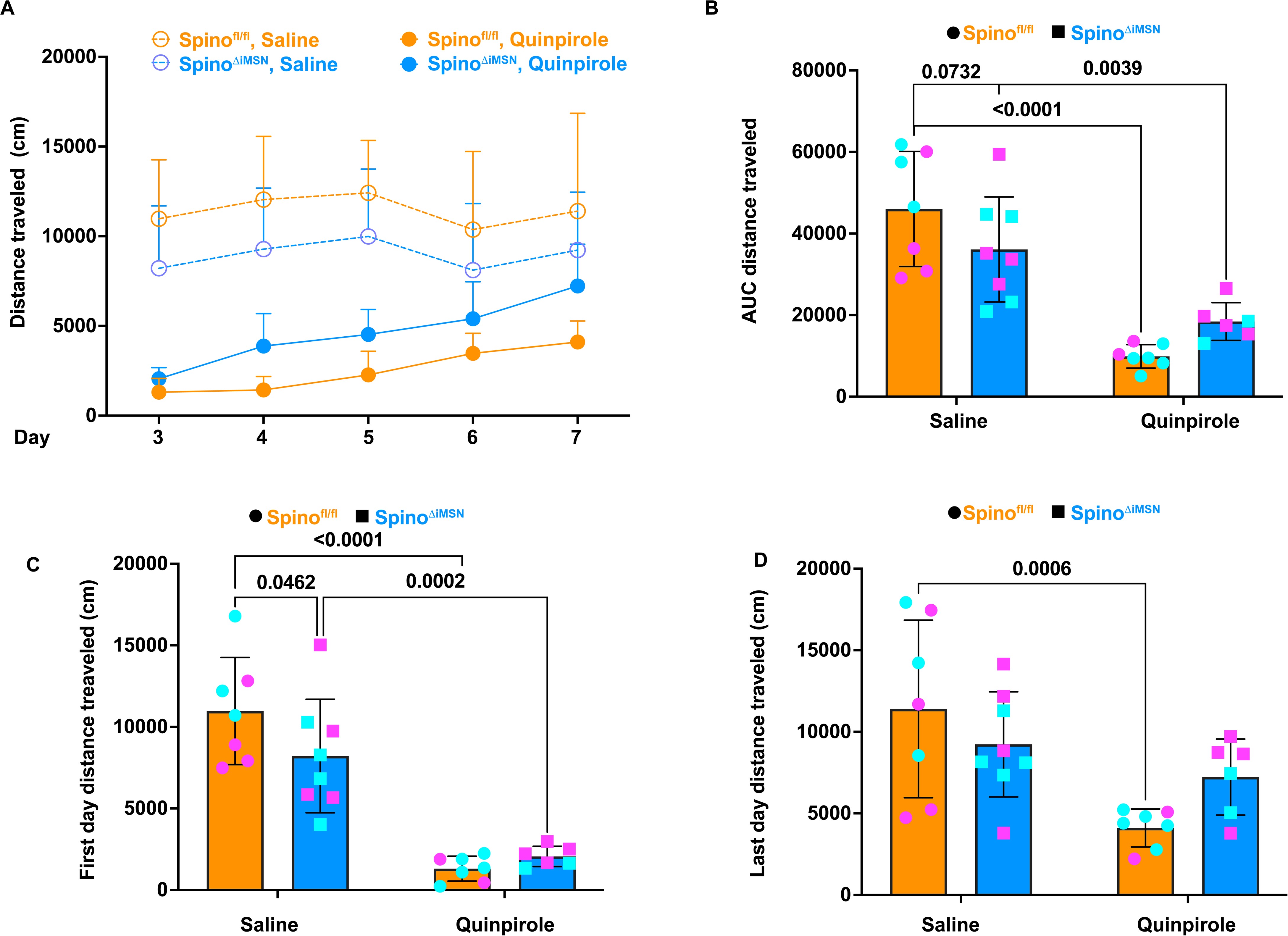
Loss of spinophilin conditionally from iMSNs did not abrogate the locomotor suppression of quinpirole. Spino^fl/fl^ or Spino^ΔiMSN^, 2–8-month-old male or female mice were intraperitoneally injected with saline daily for 2 days. Subsequently, mice were injected with saline or quinpirole daily for 5 days. The total distance traveled was recorded over 1 hour using the open-field test. **A.** 3-way ANOVA analysis revealed significant treatment (p<.0001), day (p<.0004), treatment x day (p=.0004), and treatment x genotype (p=.0257) effects **B.** AUC of distance traveled for each animal was calculated and a 2-way ANOVA, revealed significant treatment (p<.0001) and treatment x genotype (p=.0258) effects. An uncorrected Fisher’s least significant difference (LSD) showed significantly decreased movement in quinpirole treated Spino^fl/fl^ and Spino^ΔiMSN^ mice compared to their corresponding saline controls (p<.0001 and p=.0039, respectively) **C.** A 2-way ANOVA on the distance traveled on the first day was performed. There was a significant treatment effect (p<.0001). A post hoc fisher’s LSD found a significant movement decrease in quinpirole treated Spino^fl/fl^ and Spino^ΔiMSN^ quinpirole treated animals compared to their corresponding saline controls (p<.0001, p=.0002, respectively). Morever, there was a significant decrease in movement in Spino^ΔiMSN^ saline-treated mice compared with saline-treated Spino^fl/fl^ mice at day 1 (p=0.0462). **E)** A 2-way ANOVA on the distance traveled on the last day was performed. There was a significant treatment effect (p=0.0016). A post hoc fisher’s LSD found a significant movement decrease (p=.0006) only in the quinpirole treated Spino^fl/fl^ group compared to their saline controls. N = 6-8 per group. Mean + (A) or +/-standard deviation are shown. Magenta colors represent female mice, cyan colors represent male mice.

We calculated the AUC for each animal and analyzed the overall distance traveled across the groups (**Figure 6B**). A 2-way ANOVA revealed treatment (p<0.0001) and treatment x genotype (p=0.0258) effects. The post hoc analysis from uncorrected Fisher’s LSD demonstrated the decrease in movement in quinpirole-treated Spino^fl/fl^ mice compared to Spino^fl/fl^ saline-treated mice (p<0.0001). Additionally, the post hoc analysis demonstrated the significant decrease in movement of Spino^ΔiMSNs^ treated quinpirole compared to saline treated (p=0.0039) (**Figure 6B**). However, unlike in the global Spino KOs, we did not observe a significant difference in the AUC between the quinpirole-treated Spino^fl/fl^ and Spino^ΔiMSN^ mice (p=0.1458).

We also compared within the first and last injection days. For the first day, there was a significant treatment effect (p<.0001). Using an uncorrected Fisher’s LSD on the first day of treatment, there was a significant reduction in distance traveled between the quinpirole compared to saline treatment in either Spino^fl/fl^ or Spino^ΔiMSN^ animals (p<.0001, p=0.0002, respectively) There was a significant reduction of movement in saline treated Spino^ΔiMSN^ and Spino^fl/fl^ mice (p=0.0462) **(Figure 6C).** For the last day (**Figure 6D**), there was an overall treatment effect (p=0.0016). For the multiple comparisons test, there was a significant reduction in distance traveled between the quinpirole compared to saline treatment in Spino^fl/fl^ animals (p=.0006). However, there was no difference between the distance traveled in the quinpirole compared to saline treatment in the Spino^ΔiMSN^ mice. Therefore, overall, loss of spinophilin in iMSNs attenuated locomotor suppression caused by quinpirole, but did not fully recapitulate the effects observed in the global knockout mice. Therefore, we wanted to determine how loss of spinophilin in other striatal, D2R-containing neurons affected quinpirole-induced changes in locomotion.

### Spinophilin in cholinergic interneurons (CINs) contributes to quinpirole-induced locomotor suppression

While CINs only make up a few percent of all striatal neurons, they have major impacts on striatal function. Moreover, eighty percent of dopamine receptors on the surface of CINs are D2Rs and D2Rs in CINs impact neuronal activity and locomotion^36–40^. Therefore, we wanted to determine if loss of spinophilin from CINs would impact quinpirole-induced locomotor suppression. We crossed mice expressing cre-recombinase under control of the choline acetyltransferase reporter (Chat-Cre) with Spino^fl/fl^ mice to create Spino^ΔCIN^ mice (see methods). Spino^fl/fl^ or Spino^ΔCIN^ mice were injected with saline or quinpirole as described above. Locomotion was measured (**Figure 7A and S8**). Using a 3-way ANOVA, we found significant day (p<0.0001), treatment (p<0.0001), and day x treatment (p=0.0003) effects. We also conducted a 2-way ANOVA analysis among the four groups to further analyze the significant variable interactions. When comparing the treatment of saline and quinpirole in Spino^ΔCIN^ mice, we found significant day (p=.0004), treatment (p=0.0025), and day x treatment (p=.0225) effects. When comparing the treatment of saline and quinpirole in the Spino^fl/fl^ mice, we found day (p=0.0137) and treatment (p<0.0001) effects. Finally, we analyzed the different genotypes within the quinpirole treatment group. We observed day (p<0.0001) and genotype (p=0.0098) effects with no interaction between the two variables.

**Figure 7.**
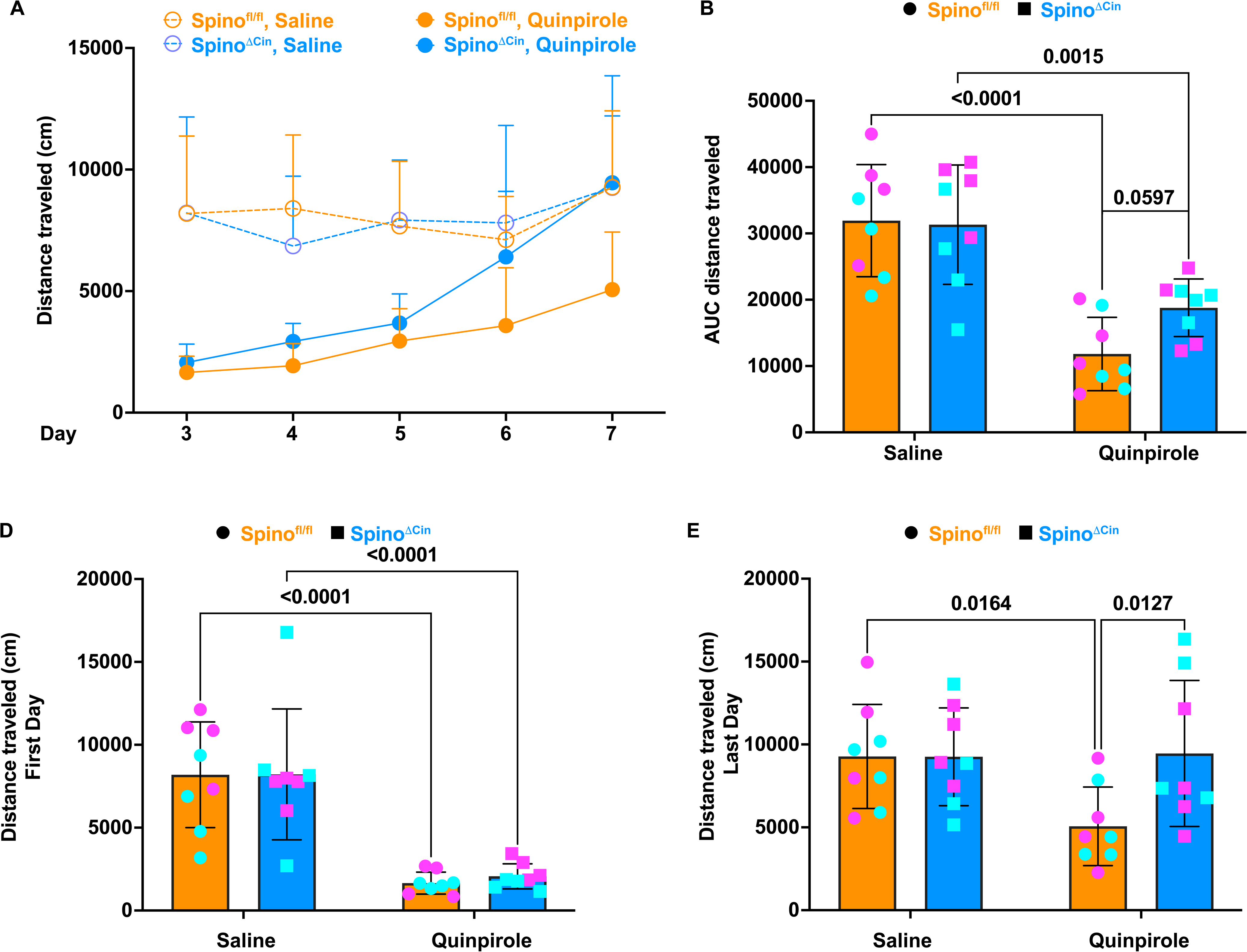
Loss of spinophilin conditionally from CINs terminated the locomotor suppression of quinpirole on the last day. Spino^fl/fl^ or Spino^ΔCIN^, 2–8-month-old male or female mice were intraperitoneally injected with saline daily for 2 days. Subsequently, mice were injected with saline or quinpirole daily for 5 days. The total distance traveled was recorded over 1 hour using the open-field test. **A.** 3-way ANOVA analysis revealed significant treatment (p<.0001), day (p<.0001), day x treatment (p=.0003) effects. **B.** AUC of distance traveled for each animal was calculated and a 2-way ANOVA, revealed a significant treatment effect (p<.0001). There was a significantly decreased movement in quinpirole-treated Spino^fl/fl^ and Spino^ΔiMSN^ mice compared to their corresponding saline controls (p<.0001, p=.0015, respectively). **C.** A 2-way ANOVA on the distance traveled on the first day was performed. There was a significant treatment effect (p<.0001). A post hoc fisher’s LSD found a significant decrease in distance traveled in the quinpirole treated Spino^fl/fl^ and Spino^ΔiMSN^ quinpirole treated animals compared to their corresponding saline controls (p<.0001, p<.0001, respectively)**. D.** A 2-way ANOVA on the distance traveled on the last day was performed. There were trending genotype (p=.0712), treatment (p=0.0963), and treatment x genotype (p=0.0689) effects. A post hoc fisher’s LSD found a decrease in distance traveled in Spino^ΔCIN^ quinpirole treated compared to saline controls (p=0.0164) and a significant increase in movement in the quinpirole treated Spino^ΔCIN^ compared with Spino^fl/fl^ quinpirole-treated mice (p=.0127). n = 8 per group. Mean + (A) or +/-standard deviation are shown. Magenta colors represent female mice, cyan colors represent male mice.

Using AUC analysis for each animal, followed by 2-way ANOVA analysis and uncorrected Fisher’s LSD (**Figure 7B**), we found a significant treatment effect (p<0.0001). There was a significant decrease in total distance traveled in both Spino^fl/fl^ and Spino^ΔCIN^ groups treated with quinpirole compared to saline (p=0.0001, p=0.0015, respectively). In addition, there was a trend for a difference between the AUC in the quinpirole-treated Spino^fl/fl^ compared with Spino^ΔCIN^ mice (p=0.0597).

We next evaluated the effects of spinophilin genotype on the initial exposure to quinpirole. On the first day, there was a significant treatment effect (p<.0001). There was a significant reduction in distance traveled between the quinpirole compared to saline treatment in both Spino^fl/fl^ and Spino^ΔCIN^ mice (p<.0001, p<0.0001, respectively) (**Figure 7C**). In contrast, when evaluating the last day of treatment, there was no overall treatment effect (p=0.3410), but trending genotype (p=0.0712), treatment (p=0.0963), and genotype x treatment (p=0.0689) effects (**Figure 7D**). Using post hoc analysis on the last day, there was a significant increase in total distance traveled in Spino^ΔCIN^ compared to Spino^fl/fl^ mice treated with quinpirole (p=0.0127). Additionally, there was a significant decrease in total distance traveled in Spino^fl/fl^ mice treated with quinpirole compared to saline (p=0.0164). Collectively, these findings suggest that while loss of spinophilin from CINs alone did not recapitulate the effect of quinpirole in Spino^-/-^ mice, loss of spinophilin from CINs enhanced locomotion in quinpirole-treated animals at the later time points.

## DISCUSSION

### Spinophilin-dependent regulation of the D2R protein interactome in Neuro2A cells

Using proximity labeling and proteomics approaches, we found that overexpression of spinophilin in Neuro2A cells enhanced the abundance of proteins that were proximal to overexpressed, UltraID-labeled D2R. When evaluating overall D2R proximity labeling, we identified multiple ribosomal proteins and ribosomal pathways. This is consistent with these experimental approaches commonly identifying proteins involved in mRNA function and translation, as these processes are required for production of these overexpressed proteins. However, when we compared the effect of WT spinophilin overexpression compared to no overexpression or overexpression of a PP1 binding-deficient mutant spinophilin, we observed greater overall protein association in the context of WT spinophilin overexpression. Intriguingly, most of the pathways associated with proteins with the greatest increase in association with the D2R are linked to the mitochondria or mitochondrial function. Additionally, within the top 50 proteins with the largest fold increase, we identified proteins associated with the intermediate filament cytoskeleton and vesicle/Golgi trafficking. The labeling radius of BioID is ∼10 nm^41^. For our specific protein, we have added 25 amino acid linker (loxp and ALFA-tag sequences) that would slightly (∼ 1 nm) enhance the distance from the D2R. Therefore, while spinophilin enhances the proximity of the D2R with cytosolic proteins such as mitochondrial and cytoskeletal proteins, it is unclear if it is increasing a direct interaction with these proteins and organelles. Overall, our data suggest that spinophilin is increasing the intracellular localization of the D2R subcellular localization, which would be predicted to limit receptor function, at least for canonical signaling via inhibition of the PKA pathway. However, as arrestin signaling may still occur, it is unclear if spinophilin impacts arrestin signaling via the D2R. Importantly, previous studies in the alpha adrenergic receptor found that spinophilin reduced arrestin actions on this receptor and reduced its arrestin-dependent internalization^22^; however, future studies need to probe spinophilin’s effects on arrestin-dependent regulation of D2R signaling.

### Mechanisms of spinophilin-dependent regulation of D2R interactome

D2Rs are localized in soma and dendrites/dendritic spines of striatal neurons^32^. Subcellularly, D2Rs are expressed on both the plasma membrane and in intracellular membranes (endomembranes) that are associated with mitochondria^32^. Moreover, quinpirole increased the association with these endomembranes in the ventral striatum^32^. Other studies have also found that treatment with agonist causes the D2R to internalize which can then be recycled or degraded^33,34^. Interestingly, our data demonstrate that overexpression of WT, but not F451A mutant, spinophilin enhanced the association of D2R with intracellular proteins, including vesicle trafficking, intermediate filament, and mitochondrial proteins. Moreover, quinpirole treatment enhanced the co-immunoprecipitation of the postsynaptically-enriched long-isoform of the D2R with actin cytoskeleton, cytosolic, and mitochondrial proteins. Conversely, loss of spinophilin limited quinpirole-induced increases in D2R interactions with these intracellular proteins. Therefore, our data suggest that spinophilin mediates quinpirole-induced intracellular trafficking of the D2R in striatal neurons. In addition to the D2R, spinophilin interacts with multiple Gαi G protein-coupled receptors, such as opioid receptors, α2 adrenergic receptors, and the M3 muscarinic receptor^19,21,22,42–44^. However, the effects of loss of spinophilin on receptor internalization are distinct depending on the receptor and complex. For instance, loss of spinophilin enhanced the internalization of the α2-adrenergic receptor, suggesting that spinophilin stabilizes the receptor at the surface^45^. Conversely, overexpression of spinophilin enhanced mu opioid receptor internalization whereas spinophilin knockout decreased this internalization^19^. Therefore, spinophilin has disparate actions on different G protein-coupled receptors and our data suggest that spinophilin increases the internalization of the D2R, similar to effects on the mu opioid receptor and opposite from actions on the α2-adrenergic receptor.

### Spinophilin limits postsynaptic D2R response to agonist

Acute activation of D2R by agonists, including quinpirole, suppresses locomotion initially, but a tolerance may develop to this locomotor suppression, depending on the dose^8,29^. However, there are dose-, time-, and species-specific effects of quinpirole^21,22^ ^29,46–48^. Therefore, we measured locomotor activity in WT C57Bl6/J mice in response to daily injections with 1 mg/kg, 3 mg/kg, or 10 mg/kg of quinpirole for 5 days. We found that 3 mg/kg of quinpirole had the most robust and consistent suppression of locomotion across the 5 days of administration. We next evaluated the effect of global loss of spinophilin on quinpirole-induced locomotor suppression to this 3 mg/kg dose of quinpirole. Loss of spinophilin had no effect on the locomotor suppressive effects of quinpirole following the first quinpirole injection. However, we found that global loss of spinophilin led to significant abrogation of locomotor suppression of quinpirole and spinophilin knockout mice were hyperlocomotive on the final quinpirole injection when compared with saline-treated, Spino^-/-^ mice. Different dopamine receptors are cell type-specifically expressed and play unique roles in dopamine signaling. Importantly, the long isoform of the D2R (D2L) is thought to exclusively be expressed postsynaptically while both the D2L and short (D2S) isoforms of the D2R are presynaptically expressed^49^. However, within presynaptic sites, the D2S and D2L have unique effects^11^. For instance, the D2S is thought to mediate the locomotor stimulating actions of psychostimulants^50^. Moreover, loss of D2S attenuated the locomotor suppression caused by quinpirole whereas loss of D2L permitted the locomotor suppressive actions of quinpirole, suggesting that the D2S is responsible for locomotor suppression^51^. Conversely, knockout of the D2L led to decreased motor activity in the home cage, suggesting D2L promotes locomotion^52^. While our initial studies in global spinophilin knockout mice found an increased tolerance to the locomotor suppressive effects of quinpirole, if this tolerance is due to decreased presynaptic or increased postsynaptic action of D2Rs is unclear. Critically, our data suggest that loss of spinophilin in striatal neurons, which receive dopamine input, is playing a role in this response as loss of spinophilin in either iMSNs or CINs led to a tolerance to the locomotor suppressive effects of quinpirole. Intriguingly, loss of spinophilin in iMSNs appeared to be responsible for earlier tolerance to the locomotive effects whereas loss in CINs was responsible for later (days 4 and 5) tolerance to the locomotor suppressive effects. Overall, while loss in either population did not fully recapitulate the effect of global knockout, it may be that the effects are additive and striatal iMSN and CIN spinophilin together are responsible for limiting postsynaptic D2R activity in the context of repeated quinpirole injections. While we cannot rule out effects of spinophilin in other D2R-containing cells such as dopamine neurons, loss of spinophilin in these neurons would be predicted to increase D2R function which would decrease neuronal activity and lead to decreased dopamine release and movement. Importantly as spinophilin was not observed in presynaptic terminals, at least in primate cortex^53,54^, we posit that spinophilin regulation of dopamine neuron D2Rs would be restricted to sites of postsynaptic D2Rs and not at presynaptic autoreceptor D2Rs. However, while less likely, we cannot rule out an effect of spinophilin on presynaptic autoreceptors as spinophilin has been shown to be presynaptic at the drosophila neuromuscular junction^55^.

### Implications of spinophilin-dependent regulation of D2R interactome in PD

According to the Parkinson’s Foundation, there will be 1.2 million Americans living with PD by 2030^56^. Dopamine replacement therapy and medications that target D2R specifically have issues with “wearing off” or dyskinesias^57^. Therefore, there is a crucial need to find alternative targets that assist in treating motor dysfunction associated with striatal disorders such as PD. Dopamine replacement therapies and D2R agonists are used to treat motor deficits associated with PD. They work because there is a postsynaptic upregulation of dopamine receptors due to constant dopamine depletion observed in PD^58^. This increase occurs early but seems to wane over time as dopamine replacement therapies and dopamine agonists are used to treat the disease^58^. Our data suggest that the agonist-induced internalization of the D2R requires spinophilin. Therefore, inhibiting spinophilin-dependent regulation of the D2R may be a potential mode to limit D2R internalization and maintain increased D2R function in the presence of D2R agonists used for treating PD-associated motor deficits. Moreover, we found PD as a major KEGG term across all of our samples with overlapping and distinct proteins and protein groups matching the PD term detected (**Figure S9**). Therefore, our studies further link spinophilin regulation of D2R to PD and future studies will need to evaluate how loss of spinophilin in iMSNs and/or CINs impacts D2R agonist efficacy in the context of a dopamine depleted state or other animal models for PD.

### Implications of spinophilin-dependent regulation of D2R interactome in dopamine neuron protection in PD

Consistent with a role for the D2R in PD, PD was one of the top 10 enriched pathways observed in the KEGG enrichment analysis across all interactome maps that were analyzed. When evaluating proteins that were modulated by spinophilin expression or quinpirole treatment that matched to the PD KEGG pathway, we discovered 3 main classes of proteins: proteins involved in oxidative phosphorylation (mitochondrial proteins), proteasomal proteins, and microtubule cytoskeleton proteins (**Figure S9**). We posit that these proteins are now in close proximity to the D2R due to its internalization; however, how D2R signaling may impact the function of these proteins is a critical future study. Importantly, mitochondrial dysfunction is a main driver of PD as genetic mutations in mitochondrial proteins are causative for early onset PD^59,60^. Moreover, D2R activation enhances mitochondrial function. For example, one study found that quinpirole ameliorated rotenone-induced mitochondrial damage in a neuroblastoma cell line expressing mutant Lrrk2, a protein associated with PD^61^. Also, quinpirole administration limited toxicity and motor deficits in mice treated with the mitochondrial toxin, MPTP^62^. In contrast to these neuroprotective effects, dopamine agonists impaired mitochondrial movement, at least in hippocampal neurons, which may impact localization of these important energy-producing organelles^63^. Given these effects of quinpirole and that spinophilin normally seems to limit D2R responsivity to quinpirole, it may be that decreasing spinophilin’s interaction with the D2R would enhance D2R activity and response to quinpirole which in turn may promote the protection of D2R-expressing nigral dopamine neurons. While current studies have focused on spinophilin in striatal neurons, it may have similar roles in regulating quinpirole-induced internalization of the D2R in dopamine neurons. Interestingly, alterations in D2Rs on dopamine neurons is required for locomotor sensitization to psychostimulants. Specifically, the D2S isoform that is expressed on midbrain dopamine neurons is sufficient to increase locomotion in responses to cocaine^64^. Previously, we and others found that global loss of spinophilin decreased locomotor sensitization caused by cocaine or amphetamine^15,31^. This is interesting as global loss of spinophilin decreased motor output in response to repeated treatments of drug rather than what is observed in our current studies where we see an increase in motor output in spinophilin knockout mice in response to repeated administration of quinpirole. Importantly, we found that knockout of spinophilin in either dopamine D1 receptor (D1R)-containing dMSNs or D2R-containing iMSNs had no effect on amphetamine-induced locomotor sensitization^30^. Therefore, spinophilin in another cell type, such as dopamine neurons, may be playing a role. Dopamine neurons express the D2S^9^ and spinophilin interacts with both D2S and D2L^12^. However, the two D2R isoforms have differential glycosylation and intracellular trafficking, so the regulation of the different isoforms by spinophilin may be different^10,65^. Moreover, as D2L is more postsynaptic and D2S is more presynaptic, it may be that there is an enrichment of D2L on dendrites and cell bodies of dopamine neurons and D2S may be more enriched at within axons and axon terminals. Therefore, future studies need to determine if loss of spinophilin in dopamine neurons enhances D2R responses to quinpirole and the mechanisms by which it does so. Moreover, if targeting the spinophilin D2R interaction in dopamine neurons can limit PD-associated toxicity needs to be assessed.

## CONCLUSIONS

We found that spinophilin enhances the association of the D2R with cytoplasmic proteins, including mitochondrial proteins. Moreover, multiple injections of quinpirole increased the association of the D2R with these same classes of intracellular proteins and this effect is attenuated by loss of spinophilin. Behaviorally, spinophilin in striatal neurons is responsible for limiting the postsynaptic D2R response to quinpirole as loss of spinophilin globally or in striatal iMSNs or CINs decreased the motor suppression induced by quinpirole. Our study suggests targeting spinophilin as a putative approach to enhance the efficacy of D2R agonists to treat PD-associated motor symptoms.

## LIMITATIONS OF THE STUDY

We focused our analysis on the role of spinophilin-dependent regulation of the D2R in quinpirole-induced locomotor effects. However, it is important to note that quinpirole is an agonist at both the D2R and the D3R and chronic treatment with quinpirole may have unique effects on D2R compared with D3R expression^46^. While the D3R is more enriched in the islands of Calleja within the ventral striatum, we cannot rule out effects of this receptor on locomotor activity. Additionally, the D3R may play an emerging role in PD pathology and treatment^66^. Intriguingly, the third intracellular loop of the D3R is similar to the D2R and therefore spinophilin may interact with this receptor as well.

Proximity labeling approaches will detect proteins in close proximity to not only mature protein but also recently synthesized proteins. Therefore, we have found that these approaches identify ribosomal proteins and mRNA translation machinery when probing overexpressed proteins in cells. Moreover, probing D2R interactions in brain lysates is limited by antibodies that are not robust and may have high off-target binding. Therefore, current pulldown approaches suffer from issues of specificity or/and selectivity. However, by probing the effect of spinophilin on the D2R interactome using both approaches, we have limited these concerns. Moreover, our data identified overlapping, spinophilin-dependent, pathways using both proximity labeling in a neuronal cell line and D2R co-immunoprecipitation in striatal lysates. However, these studies will require future biochemical validations and functional probing of specific spinophilin-dependent, D2R interactions.

## Supporting information

Key Resources

## RESOURCE AVAILABILITY

### Lead contact

Further Information and reagent/resource requests should be sent to, and will be fulfilled by, Anthony J. (A.J.) Baucum II, Ph.D.

### Materials Availability

- All generated plasmids were made from commercially available plasmids. All DNA plasmids generated in this study are available from the lead contact without restriction.
- Conditional spinophilin knockout animals are available from the lead contact with a completed materials transfer agreement.

### Data and Code Availability

- Data
  - Raw and processed data have been uploaded to MassIVE with accession MSV000098874 and cross referenced to ProteomeXchange with accession PXD067504.
- Code
  - This study did not generate any new code.
- Other items
  - Any additional information required to reanalyze the data reported in this paper is available from the lead contact upon request

## ACKNOWLEDGEMENTS

The authors acknowledge grant support from NIH - R21AA030319 (Baucum and Atwood), NIH-R33DA041876 (Baucum), Cagiantas Fellowship (Hens), John H. Edwards Fellowship (Hens) Startup support from Department of Pharmacology and Toxicology, Stark Neurosciences Research Institute, and Strategic Recruitment Initiative, Indiana University School of Medicine (Baucum).

The mass spectrometry work performed in these studies was done by the Indiana University School of Medicine Center for Proteome Analysis. Acquisition of the IUSM Center for Proteome Analysis instrumentation used for this project was provided by the Indiana University Precision Health Initiative. The proteomics work was supported, in part, by the Indiana Clinical and Translational Sciences Institute (UL1TR002529 from the National Institutes of Health, National Center for Advancing Translational Sciences, Clinical and Translational Sciences Award) and the P30 IU Simon Comprehensive Cancer Center Support Grant (P30CA082709).

## AUTHOR CONTRIBUTIONS

Designed experiments (all authors). Technical training and/or conceptual input (all authors). Wrote paper (all authors). Performed experiments and/or analysis (all authors). Reviewed and approved manuscript (all authors).

## STAR METHODS

### EXPERIMENTAL MODEL AND STUDY PARTICIPANT DETAILS

#### Animals

Animals were maintained on a 7AM:7PM:7AM light dark cycle. Global spinophilin knockout (Spino^-/-^) mice were generated from crossing heterozygous spinophilin mice (Spino^+/-^) to generate wildtype (Spino^+/+^) and Spino^-/-^ littermates^30,31,67^. Mice with a floxed spinophilin allele (Spino^fl/fl^ or Spino ^fl/+^) and mice with spinophilin knocked out of iMSNs (Spino^ΔiMSN^) were previously described^23,30^. To generate mice with spinophilin lost in medium spiny neurons, Spino^fl/fl^ or Spino^fl/+^ mice were crossed with mice expressing Cre-recombinase under control of the Adenosine A2A (A2A) promoter (A2A-Cre and Spino^fl/fl^ or Spino^fl/+^). Spino^fl/fl^/A2A-Cre (Spino^ΔiMSN^) mice were crossed with Spino^fl/fl^ mice to generate Spino^fl/fl^ control and Spino^ΔiMSN^ mice. To generate mice with spinophilin lost in cholinergic interneurons (CINs), Spino^fl/fl^ or Spino^fl/+^ mice were crossed with mice expressing Cre-recombinase under control of the endogenous choline acetyl transferase gene (Chat-Cre and Spino^fl/fl^ or Spino^fl/+^). Spino^fl/fl^ Chat-Cre (Spino^ΔCIN^) mice were crossed with Spino^fl/fl^ mice to generate Spino^fl/fl^ control and Spino^ΔCIN^ mice. Spino^fl/fl^ control littermates were used as control lines for the Spino^ΔiMSN^ and Spino^ΔCIN^ mice. All animal procedures were performed on 2–8-month-old male and female mice between 11 AM and 6 PM in accordance with the School of Medicine Institutional Animal Care and Use Committees at IU-Indianapolis (Protocol #22024, 25030). Studies were not powered to detect sex differences due to a lack of observed sex effects in the current cohorts.

#### Cell Lines

Neuro2A (N2A) cells were purchased from ATCC and were split into passages 5 and 6 and frozen samples for long-term storage. Cell lines were not authenticated nor tested for mycoplasma. However, cells were replaced if any contamination was observed.

## METHOD DETAILS

### DNA Plasmids

All constructs used for generation of the DNA used in these studies or experiments are given in the **Key Resources Table**. We used previously described approaches to generate the D2R-Alfa-UltraID (D2R-uID) construct^17,31^. Specifically, we amplified the long-isoform of the human *Drd2* receptor from the Drd2-Tango plasmid a gift from Bryan Roth (Addgene plasmid # 66269)^68^ using specific forward (CTCTCCACAGGTGTCCAGGCGGCCGCCATGGTGATGAAGACGATCATCGCCC) and reverse primers (GCAACTAGAAGGCACAGTTATTATAACTTCGTATAGCATACATTATACGAAGTTAT ACAGTGGAGAATCTTCAGAAATG). The amplified DNA was inserted into our spinophilin-UltraID plasmid^23^ that had been digested with NotI and BamHI. Clones from the ligation were sent to Plasmidsaurus (Eugene, OR) for sequencing using Oxford Nanopore Technology. The validated GenBank sequencing file for the DNA used is given in **Data S1**.

### Transfection, biotinylation, and streptavidin pulldown of Neuro2A (N2A) cells

Neuro2A (N2A) cells were thawed and allowed to recover for four days in N2A complete media (15% FBS, 1% penicillin/ streptomycin, 1% L-glutamine, 1% sodium pyruvate, 1% NEAA). On the fourth day, the media was changed. On the seventh day, cells were passaged into a T-75 flask for maintenance and allowed to grow until the next passage in a tissue culture incubator at 37°C and 5% CO_2_. For the biotinylation assay, cells were plated in a 6-well plate at 500,000 cells per well. On the third day, cells were transfected using Lipofectamine 3000 using 5 µg of DNA following the manufacturer’s protocol. The next day, cell medium was aspirated off and cells were washed 3x with Buffer B (1X phosphate buffered saline (PBS), 0.5 mM calcium chloride, 0.5 mM magnesium chloride). Next, cells were incubated in biotin (50 μM) diluted in Buffer B for 3 to 4 hours at 37°C. Subsequently, cells were washed 2 times in buffer B, and then lysed and homogenized in 700-750 mL of RIPA buffer. Following lysis, cells were sonicated (Model FB505 sonic dismembrator; Thermo Fisher Scientific, Waltham, MA) and incubated for 5 minutes with rotating at 4°C and subsequently centrifuged at 14,000 x g for 10 minutes. A portion, 45 µl, of supernatant was mixed with 4x SDS sample buffer to generate inputs. Lysate (400 - 600 µl) was mixed with 40 µl of trypsin-resistant streptavidin (ReSyn Biosciences, Gauteng, South Africa) or neutravidin (Thermo Fisher Scientific) beads. Mixtures were incubated at 4°C with mixing overnight. The next day, the beads were washed 3 times using a magnet with IP wash buffer (150 mM NaCl, 50 mM Tris-HCl, 0.5% Triton X-100). For trypsin-resistant streptavidin, the lysates were split. For blotting, 25% of the bead mixture was centrifuged at 2000 x g for 1 minute and resuspended in 30 μL of 2x SDS sample buffer. For proteomics, 75% of the bead mixture was washed two additional times in PBS for proteomics analysis preparation. For PBS washes, beads were washed by centrifugation due to the buffer change interfering with the magnetic properties of the beads. Samples were submitted to the core in PBS for tandem mass tag (TMT) proteomics. For neutravidin pulldowns, beads were washed 3 times in IP wash buffer and then resuspended in 60 µl of 2X SDS sample buffer for immunoblotting.

### Animal Behavior

Open-field locomotion was recorded in Phenotyper Cages (Nodulus Information Technology, Wageningen, Netherlands). For dose response studies, animals were intraperitoneally (IP) injected either with saline (Teknova, S5812) as a vehicle or quinpirole (Q102 Millipore-Sigma, Burlington, MA) dissolved in saline at a volume of 10 ml/kg. Animal behavior was recorded for one hour post injection and distance traveled was quantified from video recordings using Noldus XT software. For quinpirole studies across different genotypes, all mice were injected daily for 2 days with saline followed by 5 daily injections of either saline or 3 mg/kg quinpirole. Video recordings and distance traveled for 1 hour were measured after each injection. Each day, the animal was placed in a different Phenotyper arena to minimize potential arena differences.

### Striatal Tissue Homogenization

WT and Spino^-/-^ animals which underwent behavior for 7 consecutive days were killed an hour after the last injection. Whole striatal tissues were isolated and homogenized in a 1 mL of low ionic buffer as explained previously^13,15^. 500 μL of lysates were spiked with 2 ug of D2R-L antibody and allowed to rotate at 4°C for a day. The next day, 20 μL of protein G magnetic beads (ThermoFisher Scientific) were added to each sample and allowed to rock at 4°C for 2 hours. Samples were washed with IP wash buffer 3 times followed by PBS washes for 2 times. The Beads were suspended in 250 μL of PBS and submitted to the proteomics core for analysis

### Proteomics Analysis

#### On bead digests

After washing, beads were covered with 8 M Urea, 100mM Tris hydrochloride, pH 8.5, reduced with 5 mM tris (2-carboxyethyl) phosphine hydrochloride (TCEP, #C4706, Sigma-Aldrich, St. Louis, MO) for 30 minutes at room temperature to reduce the disulfide bonds. The resulting free cysteine thiols were alkylated using 10 mM chloroacetamide (CAA, # C0267 Sigma Aldrich) for 30 minutes at RT, protected from light. Samples were diluted to 2 M Urea with 50 mM Tris pH 8.5 and proteolytic digestion was carried out with Trypsin/LysC Gold (0.4 µg, Mass Spectrometry grade, Promega Corporation V5072) overnight at 35 °C. After digestion, samples were desalted on Pierce C18 spin columns (#89870, ThermoFisher Scientific) with a wash of 0.5% TFA followed by elution in 70% acetonitrile 0.1% formic acid (FA).

#### TMTpro labeling

Samples were resuspended in 100 mM triethylammonium bicarbonate (TEAB, pH 8.5 from 1 M stock). Each sample was then labeled overnight at room temperature, with 0.5 mg of Tandem Mass Tag Pro (TMTpro™) reagent (16-plex kit, manufacturer’s instructions, #44520, lot no. lot ZA382394 for N2A UltraID pulldowns and lots YL381333 and YD370302 for striatal Immunoprecipitations; see labeling scheme below. ThermoFisher Scientific). After ensuring >90% labeling efficiency, reactions were quenched with 0.3% hydroxylamine (v/v) at room temperature for 15 minutes. Labeled peptides were then mixed and dried in a speed vacuum. A Seppak C18 column (#WAT054955, Waters, Milford, MA) was used to desalt the TMT labeled multiplex with a wash of 5% ACN, 0.1% FA and elution in 70% ACN, 0.1% FA.

### LC-MS

Approximately 4% of each TMTpro mix was analyzed using an EASY-nLC 1200 HPLC system (SCR: 014993, ThermoFisher Scientific) coupled to Eclipse™ mass spectrometer with FAIMSpro interface (ThermoFisher Scientific). Each multiplex was run on a 25 cm Aurora Ultimate column (#AUR3-25075C18, Ion Opticks, Collingwood, Victoria Australia) in a 50°C column oven with a 180-minute and 130-minute gradient. For each fraction, 2% of the sample was loaded and run at 350 nl/min with a gradient of 8-38% B over 98 minutes; 30-80% B over 10 mins; held at 80% for 2 minutes; and dropping from 80-4% B over the final 5 min (Mobile phases A: 0.1% formic acid (FA), water; B: 0.1% FA, 80% Acetonitrile. The mass spectrometer was operated in positive ion mode, default charge state of 2, advanced peak determination on, and EasyIC on. Two sets of three FAIMS CVs were utilized (-45 CV; -55 CV; -65CV in one run and -40CV; -50CV; -60CV in another) each with a cycle time of 1 s and with identical MS and MS2 parameters. Precursor scans (m/z 400-1600) were done with an orbitrap resolution of 120000, RF lens% 30, 50 ms maximum inject time, standard automatic gain control (AGC) target, minimum MS2 intensity threshold of 2.5e4, MIPS mode to peptide, including charges of 2 to 6 for fragmentation with 60 sec dynamic exclusion shared across the cycles excluding isotopes. MS2 scans were performed with a quadrupole isolation window of 0.7 m/z, 34% HCD collision energy, 50000 resolution, 200% AGC target, dynamic maximum IT, fixed first mass of 100 m/z.

#### Data analysis

Resulting RAW files were analyzed in Proteome Discover™ 2.5.0.400 (Thermo Fisher Scientific)^69^ with a *Mus musculus* UniProt reference proteome FASTA (downloaded 051322) plus common laboratory contaminants (73 sequences), spinophilin or mutant spinophilin SEQUEST HT searches were conducted with full trypsin digest, 3 maximum number missed cleavages; precursor mass tolerance of 10 ppm; and a fragment mass tolerance of 0.02 Da. Static modifications used for the search were: 1) TMTpro label on peptide N-termini, 2) TMTpro label on lysine (K) and 3) carbamidomethylation on cysteine (C) residues. Dynamic modifications used for the search were 1) oxidation on M; 2) phosphorylation on S, T or Y; 3) deamidation on N or Q; 4) acetylation on protein N-termini, 5) methionine loss on protein N-termini or 6) acetylation with methionine loss on protein N-termini. A maximum of 3 dynamic modifications were allowed per peptide. Percolator False Discovery Rate was set to a strict setting of 0.01 and a relaxed setting of 0.05. IMP-ptm-RS node was used for all modification site localization scores. Values from both unique and razor peptides were used for quantification. In the consensus workflow, peptides were not normalized or scaled; impurity value corrections were applied from the TMTpro lot information. Reporter ion quantification was performed using S/N values with minimum quality control filters set to an average S/N threshold of 5 and co-isolation threshold of 30%.

##### Labeling scheme – N2A UltraID pulldowns

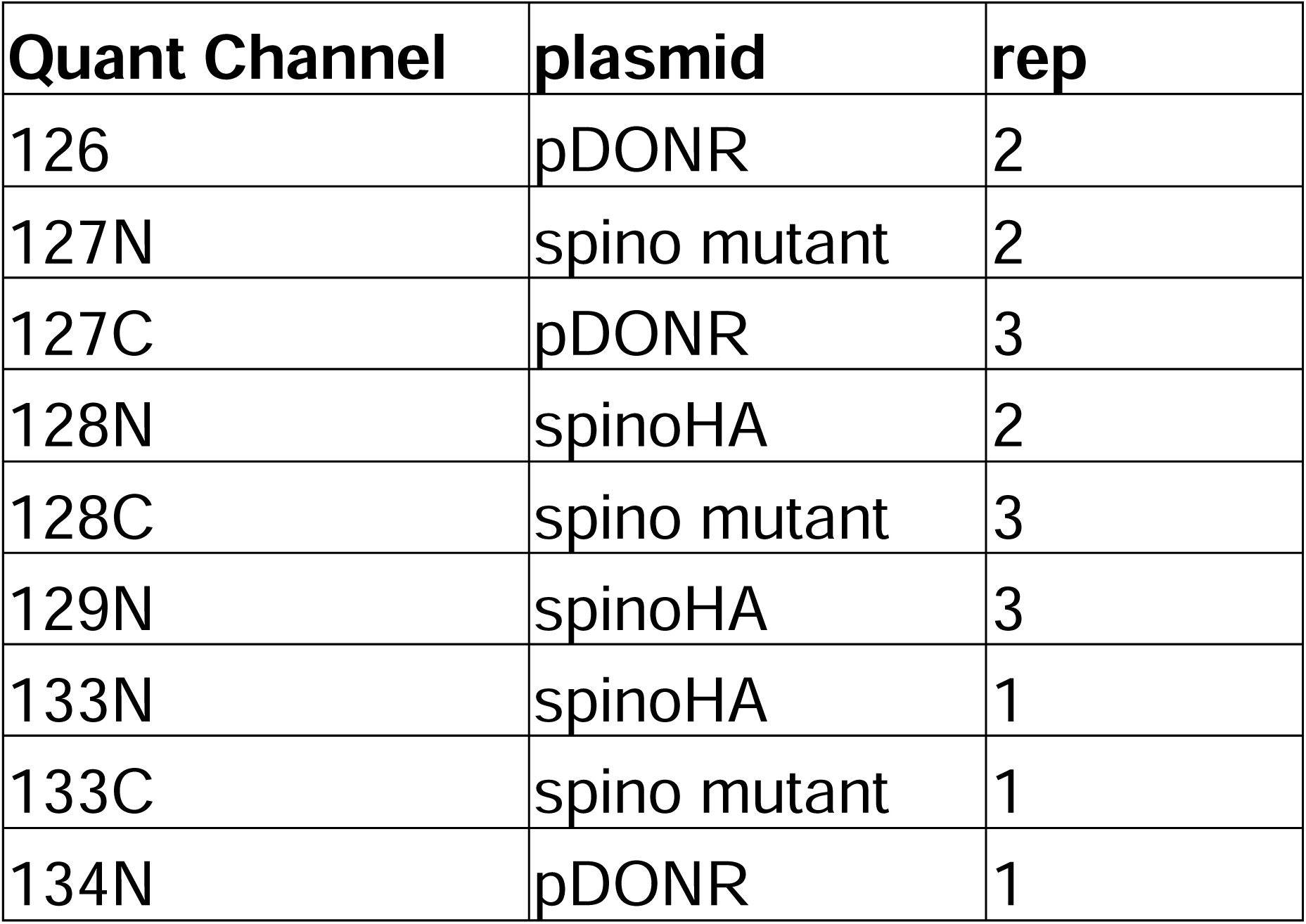

**Labeling scheme – N2A UltraID pulldowns.**
| Quant Channel | plasmid | rep |
| --- | --- | --- |
| 126 | pDONR | 2 |
| 127N | spino mutant | 2 |
| 127C | pDONR | 3 |
| 128N | spinoHA | 2 |
| 128C | spino mutant | 3 |
| 129N | spinoHA | 3 |
| 133N | spinoHA | 1 |
| 133C | spino mutant | 1 |
| 134N | pDONR | 1 |

##### Labeling scheme ȓ Striatal Immunoprecipitates

**Labeling scheme – Striatal Immunoprecipitates.**
| Quan Channel | ID | sex | genotype | treatment |
| --- | --- | --- | --- | --- |
| 126 | 12306 | M | WT | saline |
| 127N | 12301 | F | WT | saline |
| 127C | 12341 | M | WT | quinpirole |
| 128N | 12343 | M | WT | quinpirole |
| 128C | 12336 | F | WT | quinpirole |
| 133N | 12385 | F | KO | saline |
| 133C | 12388 | F | KO | saline |
| 134N | 12342 | M | KO | saline |
| 134C | 12383 | F | KO | quinpirole |
| 135N | 12300 | F | KO | quinpirole |

#### Immunoblotting

Studies were performed like our previous publication^23^. Specifically, beads were heated at 70°C for 10 minutes and then placed on a magnet (trypsin-resistant streptavidin beads) or centrifuged (neutravidin beads) at 2000 x g for 1 minutes. Pulldown sample (30 µl) was loaded onto a Criterion Gel (Bio-Rad Laboratories, Hercules, CA). Samples were electrophoresed and subsequently transferred to nitrocellulose membrane using a Trans-blot Turbo (Bio-Rad Laboratories) using a 30-minute, standard semi-dry transfer setting. Blots were imaged using an Odyssey M (LI-COR Biosciences, Lincoln, NE) Infrared imager and Image Studio was used to quantify fluorescence intensity and export the images.

## QUANTIFICATION AND STATISTICAL ANALYSES

### Enrichment analyses from STRING-db

Enrichment analysis using gene ontology (GO) and Kyoto Encyclopedia of Genes and Genomes (KEGG) was performed using STRING-db^25–28^. The full enrichment lists as well as FDR and strength for the Neuro2A and striatal lysate pulldowns are found in **Tables S4, S5, S6, S9, S11, and S12.**

### TMT proteomics

P-values (uncorrected) from t-tests comparing protein expression across two groups (WT spinophilin/non-transfected controls, F451A spinophilin/non-transfected controls, or F451A/WT spinophilin) are given in Table S3. P-values from t-tests comparing protein expression across two groups (Spino^+/+^ quinpirole/Spino^+/+^ Saline or Quinpirole Spino^-/-^/Quinpirole Spino^+/+^) are given in **Table S10**.

### Behavioral Analyses

P-values for generating volcano plots from proteomics data were calculated in Excel (Microsoft, Redmond, WA) using protein abundance based t-test across two comparison subgroups (individual protein). Statistical analyses for behaviorall assays were performed using Prism (GraphPad, San Diego, CA. Version 10) using 2-way or 3-way ANOVA with tests. A 2-way ANOVA followed by Dunnet’s multiple comparisons test was used to analyze the dose response curve. An area under the curve (AUC) was calculated for each individual animal and the AUCs across groups were analyzed with a 2-way ANOVA and uncorrected Fisher’s LSD post hoc test. The first and last day of behavior were analyzed using a 2-way ANOVA analysis and followed by uncorrected Fisher’s LSD post hoc analysis. Graphs display mean with standard deviation. All statistical analyses for behavioral analysis are given in **Table S13A-M**. The following tabs correspond to the following figure panels.

| Table Panel | Figure Panel |
| --- | --- |
| S13A | S3 |
| S13B | 4B |
| S13C | 4C |
| S13D | 4D |
| S13E | 4E |
| S13F | 6A |
| S13G | 6B |
| S13H | 6C |
| S13I | 6D |
| S13J | 7A |
| S13K | 7B |
| S13L | 7C |
| S13M | 7D |

## SUPPLEMENTAL TABLE LEGENDS (Non-PDF Format)

**Table S1. List of all proteins and their abundances detected in the D2R-ultraID (D2R-uID) pulldown in neuro2A cells.** Neuro2A cells were transfected with D2R-uID with iCre, D2R-uID with WT spinophilin and iCre, or D2R-uID with F451A mutant spinophilin and iCre. Trypsin-resistant streptavidin pulldowns were subjected to tryptic digestion and tandem-mass tag labeling followed by mass-spectrometry. The detected peptides were searched against the mouse database with human HA-spinophilin and D2R-uID appended. A total of 1,785 proteins were detected.

**Table S2. List of all peptides and their abundances detected in the D2R-uID pulldown in neuro2A cells.** Neuro2A cells were transfected with D2R-uID with iCre, D2R-uID with WT spinophilin and iCre, or D2R-uID with F451A mutant spinophilin and iCre. Trypsin-resistant streptavidin pulldowns were subjected to tryptic digestion and tandem-mass tag labeling followed by mass-spectrometry. The detected peptides were searched against the mouse database with human HA-spinophilin and D2R-uID appended. A total of 11,012 peptides were detected.

**Table S3. Protein abundances across all samples normalized to D2R-uID expression.** The protein list from Table S1 was first filtered for only those proteins that had TMT abundance calculations across all samples. Each individual protein expression was normalized to D2R-uID abundance. **A.** Total normalized protein abundance list of all proteins detected across all samples. Includes contaminant proteins. Normalized ratio comparisons across paired groups are given. A total of 1,234 proteins were detected. **B.** Total normalized protein abundance list of all proteins detected across all samples with contaminant and proteins with fewer than 4 peptide spectral matches filtered out. Normalized ratio comparisons and corresponding p-value calculations are given. A total of 956 proteins were detected.

**Table S4. Pathway analysis of the Neuro2A cell D2R interactome.** Proteins from the neuro2A interactome detected across all samples, with contaminants removed, and with 4 or more peptide spectral matches were input into the STRING-db for pathway analysis. All enrichments for gene ontology for **A)** cellular component, **B)** molecular function, and **C)** biological process, and **D)** Kyoto Encyclopedia for Genes and Genomes (KEGG) pathways are listed.

**Table S5. Pathway analysis of the effects of wildtype (WT) spinophilin overexpression on the D2R interactome.** Proteins from the neuro2A interactome with a log_2_-fold change > 1 in samples with WT spinophilin overexpressed compared with no spinophilin overexpressed were input into the STRING-db for pathway analysis. All enrichments for gene ontology for **A)** cellular component, **B)** molecular function, and **C)** biological process, and **D)** Kyoto Encyclopedia for Genes and Genomes (KEGG) pathways are listed.

**Table S6. Pathway analysis of the effects of F451A mutant spinophilin overexpression on the D2R interactome.** Proteins from the neuro2A, D2R-uID interactome with a log_2_-fold change < -1 in samples with F451A spinophilin overexpressed compared with WT spinophilin overexpressed were input into the STRING-db for pathway analysis. All enrichments for gene ontology for **A)** cellular component, **B)** molecular function, and **C)** biological process, and **D)** Kyoto Encyclopedia for Genes and Genomes (KEGG) pathways are listed.

**Table S7. List of all proteins and their abundances detected in D2R immunoprecipitations from saline and quinpirole-treated spinophilin wildtype (Spino^+/+^) and knockout (Spino^-/-^) mice.** Striatal lysates from saline- and quinpirole-treated Spino^+/+^ and Spino^-/-^ mice were immunoprecipitated for the long-isoform of the D2R. Immunoprecipitates were subjected to tryptic digestion and tandem-mass tag labeling followed by mass-spectrometry. The detected peptides were searched against the mouse database. A total of 1,912 proteins were detected.

**Table S8. List of specific proteins and their abundances detected in D2R immunoprecipitations from saline and quinpirole-treated spinophilin wildtype (Spino^+/+^) and knockout (Spino^-/-^) mice.** The protein list from Table S7 was first filtered for only those proteins that had TMT abundance calculations across all samples and at least 4 peptides spectral matches per protein.

**Table S9. Pathway analysis of the striatal D2R interactome.** The complete, triaged protein list (Table S8) from all samples was input into the STRING-db for pathway analysis. All enrichments for gene ontology for **A)** cellular component, **B)** molecular function, and **C)** biological process, and **D)** Kyoto Encyclopedia for Genes and Genomes (KEGG) pathways are listed.

**Table S10. Protein abundances and ratios across all samples normalized to D2R in D2R immunoprecipitations from saline and quinpirole-treated spinophilin wildtype (Spino^+/+^) and knockout (Spino^-/-^) mice.** The protein list from Table S8 was normalized to the abundance of the D2R. The log_2_-fold change of the abundance ratio of normalized expression between the quinpirole- and saline-treated Spino^+/+^ mice or the quinpirole-treated Spino^-/-^ compared with Spino^+/+^ mice is shown.

**Table S11. Pathway analysis of the effects of quinpirole in Spino^+/+^ mice on the D2R interactome.** Proteins from the striatal D2R interactome with a log_2_-fold change > 0.5 expression between the quinpirole- and saline-treated Spino^+/+^ mice were input into the STRING-db for pathway analysis. All enrichments for gene ontology for **A)** cellular component, **B)** molecular function, and **C)** biological process, and **D)** Kyoto Encyclopedia for Genes and Genomes (KEGG) pathways are listed.

**Table S12. Pathway analysis of the effects of spinophilin on quinpirole-induced changes in the D2R interactome.** Proteins from the striatal D2R interactome with a log_2_-fold change < 0.5 expression between Spino^-/-^ compared with Spino^+/+^, quinpirole-treated, mice were input into the STRING-db for pathway analysis. All enrichments for gene ontology for **A)** cellular component, **B)** molecular function, and **C)** biological process, and **D)** Kyoto Encyclopedia for Genes and Genomes (KEGG) pathways are listed.

**Table S13. All statistical analyses performed in the manuscript separated by figure panel. Data S1. D2R-UltraID sequence verified Genbank file.**

## REFERENCES

1. Bernheimer, H., Birkmayer, W., Hornykiewicz, O., Jellinger, K., and Seitelberger, F. (1973). Brain dopamine and the syndromes of Parkinson and Huntington. Clinical, morphological and neurochemical correlations. J Neurol Sci 20, 415–455. 10.1016/0022-510x(73)90175-5.

2. Hornykiewicz, O. (1973). Dopamine in the basal ganglia. Its role and therapeutic implications (including the clinical use of L-DOPA). Br Med Bull 29, 172–178. 10.1093/oxfordjournals.bmb.a070990.

3. Pardo-Moreno, T., Garcia-Morales, V., Suleiman-Martos, S., Rivas-Dominguez, A., Mohamed-Mohamed, H., Ramos-Rodriguez, J.J., Melguizo-Rodriguez, L., and Gonzalez-Acedo, A. (2023). Current Treatments and New, Tentative Therapies for Parkinson’s Disease. Pharmaceutics 15. 10.3390/pharmaceutics15030770.

4. Hartmann, C.J., Fliegen, S., Groiss, S.J., Wojtecki, L., and Schnitzler, A. (2019). An update on best practice of deep brain stimulation in Parkinson’s disease. Ther Adv Neurol Disord 12, 1756286419838096. 10.1177/1756286419838096.

5. Bratsos, S., Karponis, D., and Saleh, S.N. (2018). Efficacy and Safety of Deep Brain Stimulation in the Treatment of Parkinson’s Disease: A Systematic Review and Meta-analysis of Randomized Controlled Trials. Cureus 10, e3474. 10.7759/cureus.3474.

6. Bateup, H.S., Santini, E., Shen, W., Birnbaum, S., Valjent, E., Surmeier, D.J., Fisone, G., Nestler, E.J., and Greengard, P. (2010). Distinct subclasses of medium spiny neurons differentially regulate striatal motor behaviors. Proc Natl Acad Sci U S A 107, 14845–14850. 10.1073/pnas.1009874107.

7. Gardoni, F., and Bellone, C. (2015). Modulation of the glutamatergic transmission by Dopamine: a focus on Parkinson, Huntington and Addiction diseases. Front Cell Neurosci 9, 25. 10.3389/fncel.2015.00025.

8. Thompson, D., Martini, L., and Whistler, J.L. (2010). Altered ratio of D1 and D2 dopamine receptors in mouse striatum is associated with behavioral sensitization to cocaine. PLoS One 5, e11038. 10.1371/journal.pone.0011038.

9. Khan, Z.U., Mrzljak, L., Gutierrez, A., de la Calle, A., and Goldman-Rakic, P.S. (1998). Prominence of the dopamine D2 short isoform in dopaminergic pathways. Proc Natl Acad Sci U S A 95, 7731–7736. 10.1073/pnas.95.13.7731.

10. Fishburn, C.S., Elazar, Z., and Fuchs, S. (1995). Differential glycosylation and intracellular trafficking for the long and short isoforms of the D2 dopamine receptor. J Biol Chem 270, 29819–29824. 10.1074/jbc.270.50.29819.

11. Gantz, S.C., Robinson, B.G., Buck, D.C., Bunzow, J.R., Neve, R.L., Williams, J.T., and Neve, K.A. (2015). Distinct regulation of dopamine D2S and D2L autoreceptor signaling by calcium. Elife 4. 10.7554/eLife.09358.

12. Smith, F.D., Oxford, G.S., and Milgram, S.L. (1999). Association of the D2 dopamine receptor third cytoplasmic loop with spinophilin, a protein phosphatase-1-interacting protein. The Journal of biological chemistry 274, 19894–19900.

13. Solis, O., Garcia-Sanz, P., Martin, A.B., Granado, N., Sanz-Magro, A., Podlesniy, P., Trullas, R., Murer, M.G., Maldonado, R., and Moratalla, R. (2021). Behavioral sensitization and cellular responses to psychostimulants are reduced in D2R knockout mice. Addict Biol 26, e12840. 10.1111/adb.12840.

14. Allen, P.B., Zachariou, V., Svenningsson, P., Lepore, A.C., Centonze, D., Costa, C., Rossi, S., Bender, G., Chen, G., Feng, J., et al. (2006). Distinct roles for spinophilin and neurabin in dopamine-mediated plasticity. Neuroscience 140, 897–911. 10.1016/j.neuroscience.2006.02.067.

15. Areal, L.B., Hamilton, A., Martins-Silva, C., Pires, R.G.W., and Ferguson, S.S.G. (2019). Neuronal scaffolding protein spinophilin is integral for cocaine-induced behavioral sensitization and ERK1/2 activation. Mol Brain 12, 15. 10.1186/s13041-019-0434-7.

16. Morris, C.W., Watkins, D.S., Pennington, T., Doud, E.H., Qi, G., Mosley, A.L., Atwood, B.K., and Baucum, A.J. (2022). 10.1101/2022.05.24.493240.

17. Hiday, A.C., Edler, M.C., Salek, A.B., Morris, C.W., Thang, M., Rentz, T.J., Rose, K.L., Jones, L.M., and Baucum, A.J., 2nd (2017). Mechanisms and Consequences of Dopamine Depletion-Induced Attenuation of the Spinophilin/Neurofilament Medium Interaction. Neural Plast 2017, 4153076. 10.1155/2017/4153076.

18. Watkins, D.S., True, J.D., Mosley, A.L., and Baucum, A.J., 2nd (2018). Proteomic Analysis of the Spinophilin Interactome in Rodent Striatum Following Psychostimulant Sensitization. Proteomes 6. 10.3390/proteomes6040053.

19. Charlton, J.J., Allen, P.B., Psifogeorgou, K., Chakravarty, S., Gomes, I., Neve, R.L., Devi, L.A., Greengard, P., Nestler, E.J., and Zachariou, V. (2008). Multiple actions of spinophilin regulate mu opioid receptor function. Neuron 58, 238–247. 10.1016/j.neuron.2008.02.006.

20. Di Sebastiano, A.R., Fahim, S., Dunn, H.A., Walther, C., Ribeiro, F.M., Cregan, S.P., Angers, S., Schmid, S., and Ferguson, S.S. (2016). Role of Spinophilin in Group I Metabotropic Glutamate Receptor Endocytosis, Signaling, and Synaptic Plasticity. The Journal of biological chemistry 291, 17602–17615. 10.1074/jbc.M116.722355.

21. Fourla, D.D., Papakonstantinou, M.P., Vrana, S.M., and Georgoussi, Z. (2012). Selective interactions of spinophilin with the C-terminal domains of the delta- and mu-opioid receptors and G proteins differentially modulate opioid receptor signaling. Cell Signal 24, 2315–2328. 10.1016/j.cellsig.2012.08.002.

22. Wang, Q., Zhao, J., Brady, A.E., Feng, J., Allen, P.B., Lefkowitz, R.J., Greengard, P., and Limbird, L.E. (2004). Spinophilin blocks arrestin actions in vitro and in vivo at G protein-coupled receptors. Science (New York, N.Y 304, 1940–1944.

23. Claeboe, E.T., Blake, K.L., Shah, N.R., Morris, C.W., Hens, B., Atwood, B.K., Absalon, S.J., Mosley, A.L., Doud, E.H., and Baucum, A.J. (2025). Proximity labeling and orthogonal nanobody pulldown (ID-oPD) approaches to map the spinophilin interactome uncover a putative role for spinophilin in protein homeostasis. bioRxiv, 2025.2001.2023.634546. 10.1101/2025.01.23.634546.

24. Kubitz, L., Bitsch, S., Zhao, X., Schmitt, K., Deweid, L., Roehrig, A., Barazzone, E.C., Valerius, O., Kolmar, H., and Bethune, J. (2022). Engineering of ultraID, a compact and hyperactive enzyme for proximity-dependent biotinylation in living cells. Commun Biol 5, 657. 10.1038/s42003-022-03604-5.

25. Szklarczyk, D., Franceschini, A., Kuhn, M., Simonovic, M., Roth, A., Minguez, P., Doerks, T., Stark, M., Muller, J., Bork, P., et al. (2011). The STRING database in 2011: functional interaction networks of proteins, globally integrated and scored. Nucleic Acids Res 39, D561–568. 10.1093/nar/gkq973.

26. Szklarczyk, D., Franceschini, A., Wyder, S., Forslund, K., Heller, D., Huerta-Cepas, J., Simonovic, M., Roth, A., Santos, A., Tsafou, K.P., et al. (2015). STRING v10: protein-protein interaction networks, integrated over the tree of life. Nucleic Acids Res 43, D447–452. 10.1093/nar/gku1003.

27. Szklarczyk, D., Gable, A.L., Nastou, K.C., Lyon, D., Kirsch, R., Pyysalo, S., Doncheva, N.T., Legeay, M., Fang, T., Bork, P., et al. (2021). The STRING database in 2021: customizable protein-protein networks, and functional characterization of user-uploaded gene/measurement sets. Nucleic Acids Res 49, D605–D612. 10.1093/nar/gkaa1074.

28. Szklarczyk, D., Morris, J.H., Cook, H., Kuhn, M., Wyder, S., Simonovic, M., Santos, A., Doncheva, N.T., Roth, A., Bork, P., et al. (2016). The STRING database in 2017: quality-controlled protein-protein association networks, made broadly accessible. Nucleic Acids Res. 10.1093/nar/gkw937.

29. Rowlett, J.K., Mattingly, B.A., and Bardo, M.T. (1995). Repeated quinpirole treatment: locomotor activity, dopamine synthesis, and effects of selective dopamine antagonists. Synapse 20, 209–216. 10.1002/syn.890200304.

30. Morris, C.W., Watkins, D.S., Shah, N.R., Pennington, T., Hens, B., Qi, G., Doud, E.H., Mosley, A.L., Atwood, B.K., and Baucum, A.J., 2nd (2023). Spinophilin Limits Metabotropic Glutamate Receptor 5 Scaffolding to the Postsynaptic Density and Cell Type Specifically Mediates Excessive Grooming. Biological psychiatry 93, 976–988. 10.1016/j.biopsych.2022.12.008.

31. Morris, C.W., Watkins, D.S., Salek, A.B., Edler, M.C., and Baucum, A.J., 2nd (2018). The association of spinophilin with disks large-associated protein 3 (SAPAP3) is regulated by metabotropic glutamate receptor (mGluR) 5. Molecular and cellular neurosciences 90, 60–69. 10.1016/j.mcn.2018.06.001.

32. Lane, D.A., Chan, J., Fitzgerald, M.L., Kearn, C.S., Mackie, K., and Pickel, V.M. (2012). Quinpirole elicits differential in vivo changes in the pre- and postsynaptic distributions of dopamine D(2) receptors in mouse striatum: relation to cannabinoid-1 (CB(1)) receptor targeting. Psychopharmacology (Berl) 221, 101–113. 10.1007/s00213-011-2553-4.

33. Namkung, Y., Dipace, C., Javitch, J.A., and Sibley, D.R. (2009). G protein-coupled receptor kinase-mediated phosphorylation regulates post-endocytic trafficking of the D2 dopamine receptor. J Biol Chem 284, 15038–15051. 10.1074/jbc.M900388200.

34. Bartlett, S.E., Enquist, J., Hopf, F.W., Lee, J.H., Gladher, F., Kharazia, V., Waldhoer, M., Mailliard, W.S., Armstrong, R., Bonci, A., and Whistler, J.L. (2005). Dopamine responsiveness is regulated by targeted sorting of D2 receptors. Proc Natl Acad Sci U S A 102, 11521–11526. 10.1073/pnas.0502418102.

35. Guo, N., Guo, W., Kralikova, M., Jiang, M., Schieren, I., Narendran, R., Slifstein, M., Abi-Dargham, A., Laruelle, M., Javitch, J.A., and Rayport, S. (2010). Impact of D2 receptor internalization on binding affinity of neuroimaging radiotracers. Neuropsychopharmacology 35, 806–817. 10.1038/npp.2009.189.

36. Dawson, V.L., Dawson, T.M., and Wamsley, J.K. (1990). Muscarinic and dopaminergic receptor subtypes on striatal cholinergic interneurons. Brain Res Bull 25, 903–912. 10.1016/0361-9230(90)90186-4.

37. Kharkwal, G., Brami-Cherrier, K., Lizardi-Ortiz, J.E., Nelson, A.B., Ramos, M., Del Barrio, D., Sulzer, D., Kreitzer, A.C., and Borrelli, E. (2016). Parkinsonism Driven by Antipsychotics Originates from Dopaminergic Control of Striatal Cholinergic Interneurons. Neuron 91, 67–78. 10.1016/j.neuron.2016.06.014.

38. Cavallaro, J., Yeisley, J., Akdogan, B., Salazar, R.E., Floeder, J.R., Balsam, P.D., and Gallo, E.F. (2023). Dopamine D2 receptors in nucleus accumbens cholinergic interneurons increase impulsive choice. Neuropsychopharmacology 48, 1309–1317. 10.1038/s41386-023-01608-1.

39. Chancey, J.H., Kellendonk, C., Javitch, J.A., and Lovinger, D.M. (2023). Dopaminergic D2 receptor modulation of striatal cholinergic interneurons contributes to sequence learning. bioRxiv, 2023.2008.2028.554807. 10.1101/2023.08.28.554807.

40. Sato, A., Sasaoka, T., Nishijo, T., and Momiyama, T. (2014). GABAergic synaptic transmission onto striatal cholinergic interneurons in dopamine D2 receptor knock-out mice. Neuroscience 263, 138–147. 10.1016/j.neuroscience.2014.01.010.

41. Kim, D.I., Birendra, K.C., Zhu, W., Motamedchaboki, K., Doye, V., and Roux, K.J. (2014). Probing nuclear pore complex architecture with proximity-dependent biotinylation. Proc Natl Acad Sci U S A 111, E2453–2461. 10.1073/pnas.1406459111.

42. Ruiz de Azua, I., Nakajima, K., Rossi, M., Cui, Y., Jou, W., Gavrilova, O., and Wess, J. (2012). Spinophilin as a novel regulator of M3 muscarinic receptor-mediated insulin release in vitro and in vivo. FASEB J 26, 4275–4286. 10.1096/fj.12-204644.

43. Wang, Q., and Limbird, L.E. (2002). Regulated interactions of the alpha 2A adrenergic receptor with spinophilin, 14-3-3zeta, and arrestin 3. The Journal of biological chemistry 277, 50589–50596.

44. Wang, Q., and Limbird, L.E. (2007). Regulation of alpha2AR trafficking and signaling by interacting proteins. Biochem Pharmacol 73, 1135–1145.

45. Brady, A.E., Wang, Q., Colbran, R.J., Allen, P.B., Greengard, P., and Limbird, L.E. (2003). Spinophilin stabilizes cell surface expression of alpha 2B-adrenergic receptors. The Journal of biological chemistry 278, 32405–32412.

46. Akhisaroglu, M., Kurtuncu, M., Manev, H., and Uz, T. (2005). Diurnal rhythms in quinpirole-induced locomotor behaviors and striatal D2/D3 receptor levels in mice. Pharmacol Biochem Behav 80, 371–377. 10.1016/j.pbb.2004.11.016.

47. Li, S.M., Collins, G.T., Paul, N.M., Grundt, P., Newman, A.H., Xu, M., Grandy, D.K., Woods, J.H., and Katz, J.L. (2010). Yawning and locomotor behavior induced by dopamine receptor agonists in mice and rats. Behav Pharmacol 21, 171–181. 10.1097/FBP.0b013e32833a5c68.

48. Park, J., Moon, E., Lim, H.J., Kim, K., Hong, Y.R., and Lee, J.H. (2023). Changes of Locomotor Activity by Dopamine D2, D3 Agonist Quinpirole in Mice Using Home-cage Monitoring System. Clin Psychopharmacol Neurosci 21, 686–692. 10.9758/cpn.22.1016.

49. Usiello, A., Baik, J.H., Rouge-Pont, F., Picetti, R., Dierich, A., LeMeur, M., Piazza, P.V., and Borrelli, E. (2000). Distinct functions of the two isoforms of dopamine D2 receptors. Nature 408, 199–203. 10.1038/35041572.

50. De Mei, C., Ramos, M., Iitaka, C., and Borrelli, E. (2009). Getting specialized: presynaptic and postsynaptic dopamine D2 receptors. Curr Opin Pharmacol 9, 53–58. 10.1016/j.coph.2008.12.002.

51. Radl, D., Chiacchiaretta, M., Lewis, R.G., Brami-Cherrier, K., Arcuri, L., and Borrelli, E. (2018). Differential regulation of striatal motor behavior and related cellular responses by dopamine D2L and D2S isoforms. Proc Natl Acad Sci U S A 115, 198–203. 10.1073/pnas.1717194115.

52. Wang, Y., Xu, R., Sasaoka, T., Tonegawa, S., Kung, M.P., and Sankoorikal, E.B. (2000). Dopamine D2 long receptor-deficient mice display alterations in striatum-dependent functions. J Neurosci 20, 8305–8314. 10.1523/JNEUROSCI.20-22-08305.2000.

53. Muly, E.C., Allen, P., Mazloom, M., Aranbayeva, Z., Greenfield, A.T., and Greengard, P. (2004). Subcellular distribution of neurabin immunolabeling in primate prefrontal cortex: comparison with spinophilin. Cereb Cortex 14, 1398–1407.

54. Muly, E.C., Smith, Y., Allen, P., and Greengard, P. (2004). Subcellular distribution of spinophilin immunolabeling in primate prefrontal cortex: localization to and within dendritic spines. The Journal of comparative neurology 469, 185–197.

55. Muhammad, K., Reddy-Alla, S., Driller, J.H., Schreiner, D., Rey, U., Bohme, M.A., Hollmann, C., Ramesh, N., Depner, H., Lutzkendorf, J., et al. (2015). Presynaptic spinophilin tunes neurexin signalling to control active zone architecture and function. Nat Commun 6, 8362. 10.1038/ncomms9362.

56. Who has parkinson’s. https://www.parkinson.org/understanding-parkinsons/statistics#:~:text=The%20combined%20direct%20and%20indirect,up%20to%20%24100%2C000%20per%20person.

57. Parkinson’s Disease. https://www.ninds.nih.gov/health-information/disorders/parkinsons-disease.

58. Kaasinen, V., Vahlberg, T., Stoessl, A.J., Strafella, A.P., and Antonini, A. (2021). Dopamine Receptors in Parkinson’s Disease: A Meta-Analysis of Imaging Studies. Mov Disord 36, 1781–1791. 10.1002/mds.28632.

59. Henrich, M.T., Oertel, W.H., Surmeier, D.J., and Geibl, F.F. (2023). Mitochondrial dysfunction in Parkinson’s disease - a key disease hallmark with therapeutic potential. Mol Neurodegener 18, 83. 10.1186/s13024-023-00676-7.

60. Thomas, B., and Beal, M.F. (2007). Parkinson’s disease. Hum Mol Genet 16 Spec No. 2, R183–194. 10.1093/hmg/ddm159.

61. Tozzi, A., Tantucci, M., Marchi, S., Mazzocchetti, P., Morari, M., Pinton, P., Mancini, A., and Calabresi, P. (2018). Dopamine D2 receptor-mediated neuroprotection in a G2019S Lrrk2 genetic model of Parkinson’s disease. Cell Death Dis 9, 204. 10.1038/s41419-017-0221-2.

62. Tang, T.T., Bi, M.X., Diao, M.N., Zhang, X.Y., Chen, L., Xiao, X., Jiao, Q., Chen, X., Yan, C.L., Du, X.X., and Jiang, H. (2023). Quinpirole ameliorates nigral dopaminergic neuron damage in Parkinson’s disease mouse model through activating GHS-R1a/D(2)R heterodimers. Acta Pharmacol Sin 44, 1564–1575. 10.1038/s41401-023-01063-0.

63. Chen, S., Owens, G.C., and Edelman, D.B. (2008). Dopamine inhibits mitochondrial motility in hippocampal neurons. PLoS One 3, e2804. 10.1371/journal.pone.0002804.

64. Welter, M., Vallone, D., Samad, T.A., Meziane, H., Usiello, A., and Borrelli, E. (2007). Absence of dopamine D2 receptors unmasks an inhibitory control over the brain circuitries activated by cocaine. Proc Natl Acad Sci U S A 104, 6840–6845. 10.1073/pnas.0610790104.

65. Kim, S.J., Kim, M.Y., Lee, E.J., Ahn, Y.S., and Baik, J.H. (2004). Distinct regulation of internalization and mitogen-activated protein kinase activation by two isoforms of the dopamine D2 receptor. Mol Endocrinol 18, 640–652. 10.1210/me.2003-0066.

66. Yang, P., Perlmutter, J.S., Benzinger, T.L.S., Morris, J.C., and Xu, J. (2020). Dopamine D3 receptor: A neglected participant in Parkinson Disease pathogenesis and treatment? Ageing Res Rev 57, 100994. 10.1016/j.arr.2019.100994.

67. Edler, M.C., Salek, A.B., Watkins, D.S., Kaur, H., Morris, C.W., Yamamoto, B.K., and Baucum, A.J., 2nd (2018). Mechanisms Regulating the Association of Protein Phosphatase 1 with Spinophilin and Neurabin. ACS Chem Neurosci 9, 2701–2712. 10.1021/acschemneuro.8b00144.

68. Kroeze, W.K., Sassano, M.F., Huang, X.P., Lansu, K., McCorvy, J.D., Giguere, P.M., Sciaky, N., and Roth, B.L. (2015). PRESTO-Tango as an open-source resource for interrogation of the druggable human GPCRome. Nat Struct Mol Biol 22, 362–369. 10.1038/nsmb.3014.

69. Orsburn, B.C. (2021). Proteome Discoverer-A Community Enhanced Data Processing Suite for Protein Informatics. Proteomes 9. 10.3390/proteomes9010015.

70. Rossi, J., Balthasar, N., Olson, D., Scott, M., Berglund, E., Lee, C.E., Choi, M.J., Lauzon, D., Lowell, B.B., and Elmquist, J.K. (2011). Melanocortin-4 receptors expressed by cholinergic neurons regulate energy balance and glucose homeostasis. Cell Metab 13, 195–204. 10.1016/j.cmet.2011.01.010.

71. Sauer, B., and Henderson, N. (1990). Targeted insertion of exogenous DNA into the eukaryotic genome by the Cre recombinase. New Biol 2, 441–449.

