## Supplemental Legends and Figures for "Striatal spinophilin enhances D2R interaction with cytosolic proteins to mediate persistent D2R agonist-induced locomotor suppression"

**Figure S1. Validation of transfections in Neuro2A cells**. A Cre-dependent ultraID-fused (DRD2-uID) construct was transfected with improved Cre-recombinase construct (iCre) along with a Pdonr empty vector (-) or wildtype (WT) or F451A mutant (FA) spinophilin. Input lysates (I) or trypsin-resistant streptavidin pulldowns (PD) were blotted with a streptavidin conjugated to Alexa 790 or with an HA antibody.

**Figure S2. Identification of the D2R interactome in Neuro2A cells**. Neuro2A cells were transfected with a Cre-dependent ultraID-fused construct, an improved Cre-recombinase construct and either a Pdonr empty vector or wildtype or F451A mutant (FA) spinophilin. Trypsin-resistant streptavidin pulldowns (PD) were performed and beads subjected to on bead digestion and tandem mass tag protemics. All proteins with at least 3 total peptide spectral matches and that were not known contaminants were input into STRING-db. The top 10 gene ontology (GO) pathways detected in the D2R interactome for **A)** cellular component, **B)** molecular function, or **C)** biological process are shown. **D.** The top 10 pathways from the Kyoto Encyclopedia for Genes and Genomes (KEGG) analysis are shown. **E.** The top 50 most abundant proteins by total peptide spectral matches were input into the STRING-db. Some of the more enriched GO pathways are highlighted.

**Figure S3**. **Quinpirole dose response on locomotor output.** Male and female, 2–8-month-old C57Bl6/J mice (n=3 per group) were injected intraperitoneally with saline, 1 mg/kg, 3 mg/kg, or 10 mg/kg quinpirole. A two-way ANOVA revealed a day (p=.0131) and treatment effect (p=.0012). Using Dunnett's multiple comparison test, the 3 mg/kg dosage maintained a significant suppression of the locomotion across the first 4 days of the behavior compared to the saline treatment (day 1: p=.0335, day 2: p=.0086, day 3: p=.0304, day 4: p=.0007, day 5: p=.07)

**Figure S4. Distance traveled across all injection days.** The same graph from figure 4B with the 2 first days of saline acclimatization injections included.

**Figure S5. STRING-db and pathway analysis of all D2R interacting proteins.** All proteins in the D2R-pulldown that had 4 or more peptide spectral matches (PSMs) across all samples and were not known contaminants were input into the STRING-db. **A-D.** The top 10 gene ontology pathways for **A)** cellular component, **B)** molecular function, and **C)** biological process, and the top 10 **D)** Kyoto Encyclopedia for Genes and Genomes (KEGG) pathways are shown. **E.** STRING-db map of the Top 50 proteins by PSM are given. N=10 total mice.

**Figure S6. Quinpirole enhances the interaction of the D2R with intracellular proteins. A.** Abundance of the D2R from the tandem mass tag (TMT) values across all 4 groups. **B.** Volcano plot of the $\log_2$-fold change and p-values of protein abundances from the TMT values between the quinpirole-treated and saline treated Spino$^{+/+}$ animals **C-F.** The proteins that had a $\log_2$-fold change of 0.5 or greater in the quinpirole-treated compared with saline-treated wildtype mice were input into the STRING-db for pathway analysis. The top 10 gene ontology pathways for **C)** cellular component, **D)** molecular function, and **E)** biological process, and the top 10 **F)** Kyoto Encyclopedia for Genes and Genomes (KEGG) pathways are shown. **G) .** STRING-db map of the Top 50 proteins by PSM are given. N=2-3 animals per group.

**Figure S7. Distance traveled across all injection days.** The same graph from figure 6A with the 2 first days of saline acclimatization injections included.

**Figure S8. Distance traveled across all injection days.** The same graph from figure 7A with the 2 first days of saline acclimatization injections included.

Figure S1

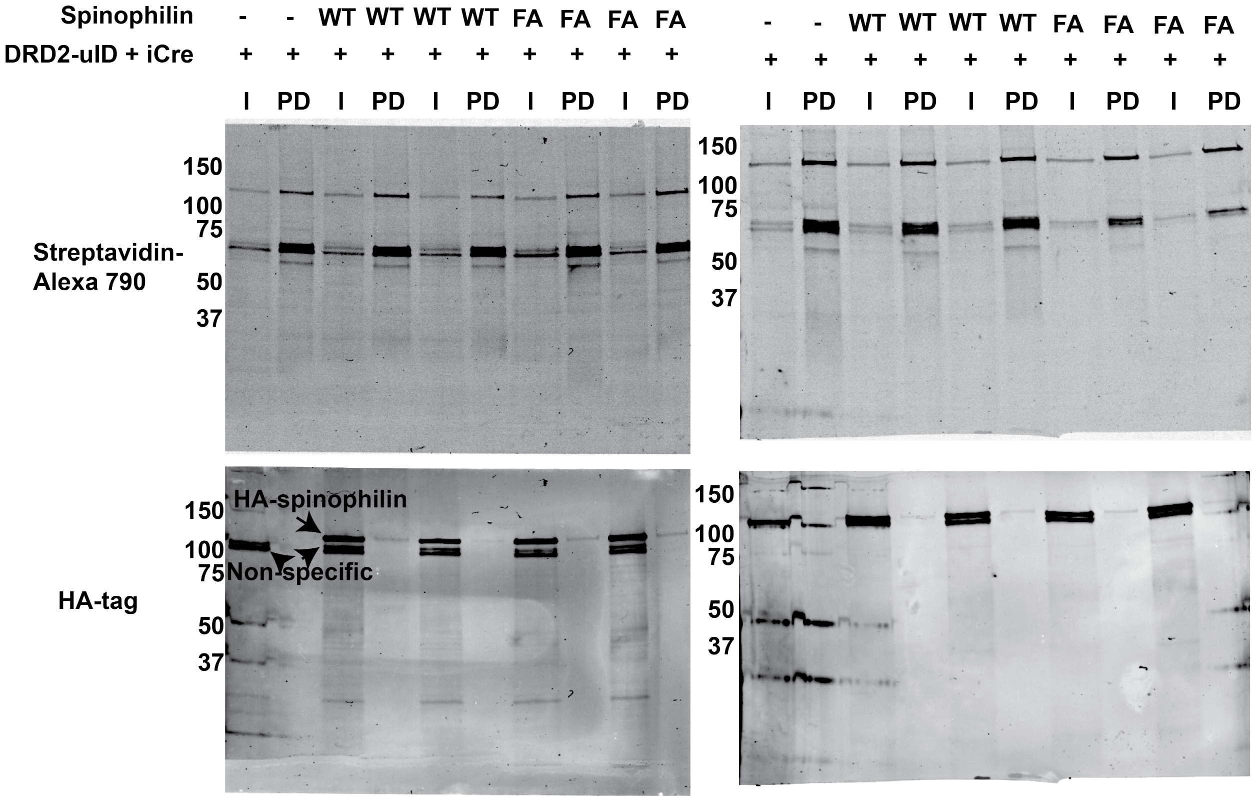
Spinophilin - - WT WT WT WT FA FA FA FA
DRD2-uID + iCre + + + + + + + + + +
I PD I PD I PD I PD I PD
Streptavidin-Alexa 790
HA-tag
150
100
75
50
37
HA-spinophilin
Non-specific
Spinophilin - - WT WT WT WT FA FA FA FA
+ + + + + + + + + +
I PD I PD I PD I PD I PD
150
100
75
50
37

Figure S2

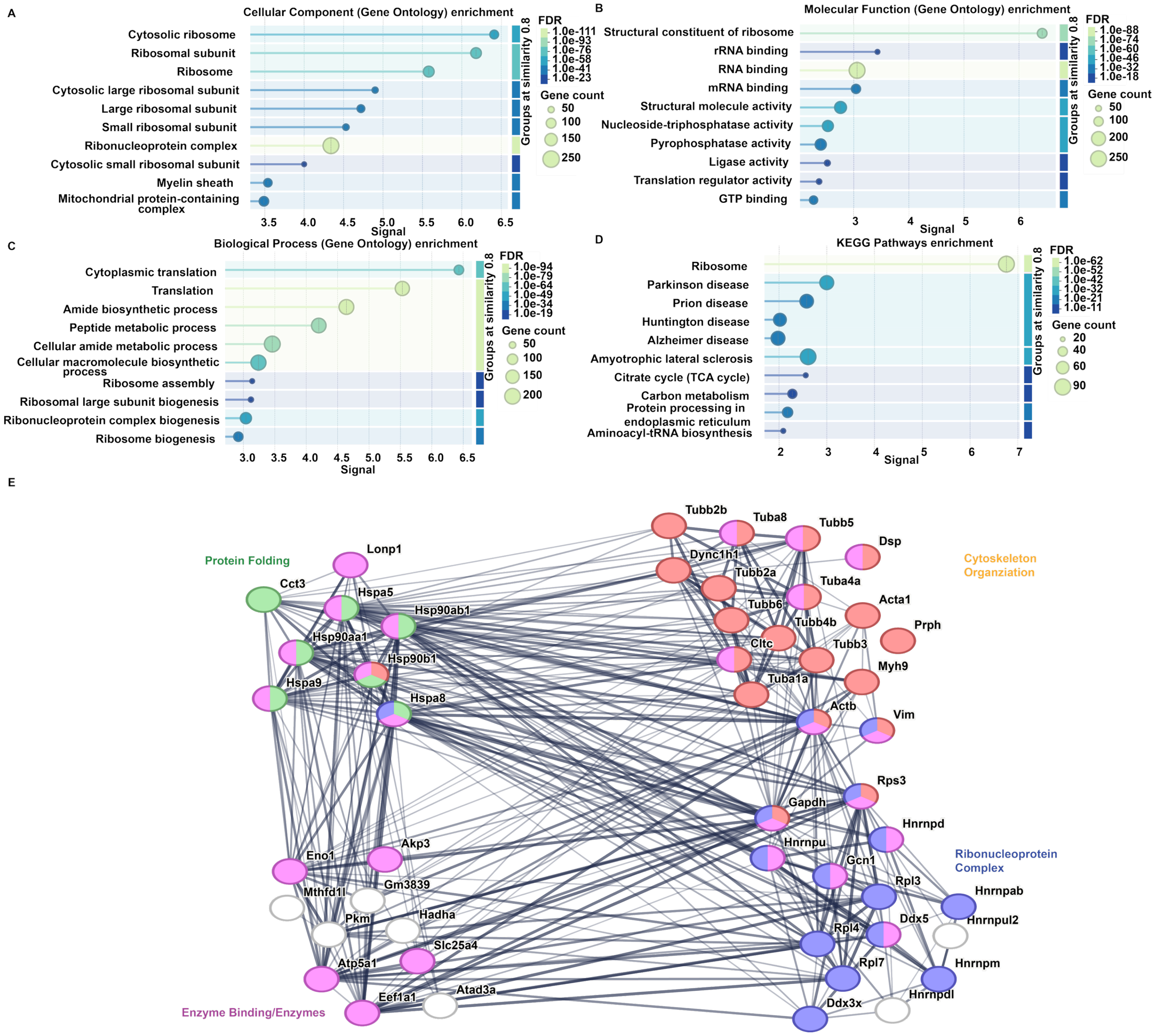
A
Cellular Component (Gene Ontology) enrichment
Cytosolic ribosome
Ribosomal subunit
Ribosome
Cytosolic large ribosomal subunit
Large ribosomal subunit
Small ribosomal subunit
Ribonucleoprotein complex
Cytosolic small ribosomal subunit
Myelin sheath
Mitochondrial protein-containing complex
Signal
FDR
Groups at similarity 0.8
Gene count
B
Molecular Function (Gene Ontology) enrichment
Structural constituent of ribosome
rRNA binding
RNA binding
mRNA binding
Structural molecule activity
Nucleoside-triphosphatase activity
Pyrophosphatase activity
Ligase activity
Translation regulator activity
GTP binding
C
Biological Process (Gene Ontology) enrichment
Cytoplasmic translation
Translation
Amide biosynthetic process
Peptide metabolic process
Cellular amide metabolic process
Cellular macromolecule biosynthetic process
Ribosome assembly
Ribosomal large subunit biogenesis
Ribonucleoprotein complex biogenesis
Ribosome biogenesis
D
KEGG Pathways enrichment
Ribosome
Parkinson disease
Prion disease
Huntington disease
Alzheimer disease
Amyotrophic lateral sclerosis
Citrate cycle (TCA cycle)
Carbon metabolism
Protein processing in endoplasmic reticulum
Aminoacyl-tRNA biosynthesis
E
Protein Folding
Cytoskeleton Organziation
Ribonucleoprotein Complex
Enzyme Binding/Enzymes
Lonp1
Cct3
Hspa5
Hsp90ab1
Hsp90aa1
Hsp90b1
Hspa9
Hspa8
Tubb2b
Tuba8
Tubb5
Dsp
Dync1h1
Tubb2a
Tuba4a
Tubb6
Acta1
Tubb4b
Prph
Cltc
Tubb3
Myh9
Tuba1a
Actb
Vim
Rps3
Gapdh
Hnrnpd
Hnrnpu
Gcn1
Rpl3
Hnrnpab
Ddx5
Hnrnpul2
Rpl4
Rpl7
Hnrnpm
Hnrnpdl
Ddx3x
Eno1
Akp3
Mthfd1l
Gm3839
Pkm
Hadha
Slc25a4
Atp5a1
Atad3a
Eef1a1

**Figure S3**

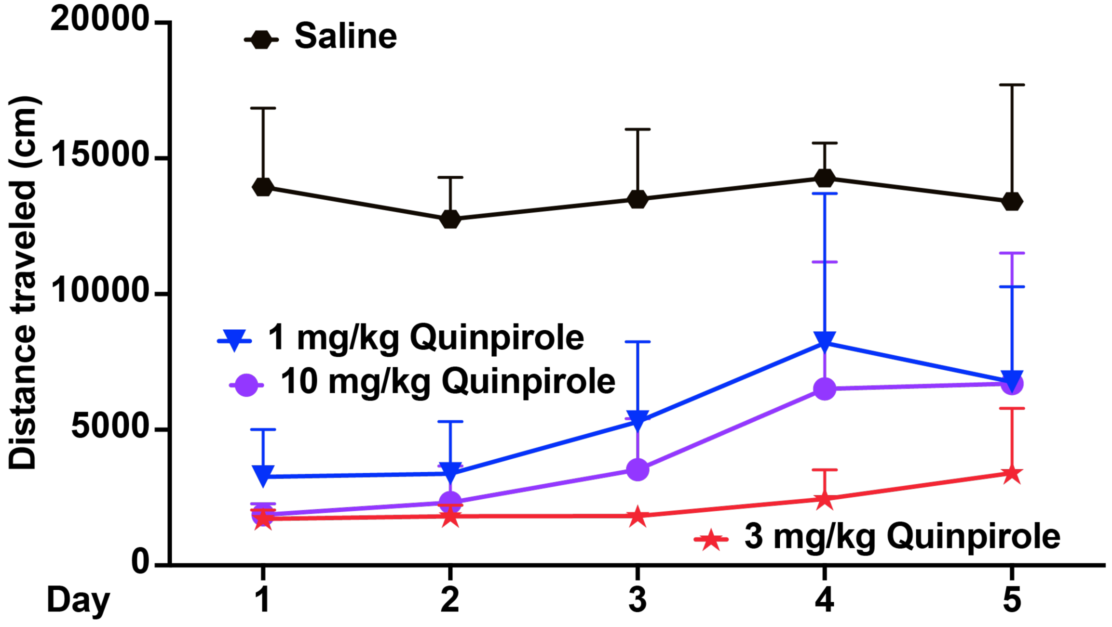

Saline
1 mg/kg Quinpirole
10 mg/kg Quinpirole
3 mg/kg Quinpirole
Distance traveled (cm)
20000
15000
10000
5000
0
Day
1
2
3
4
5

**Figure S4**

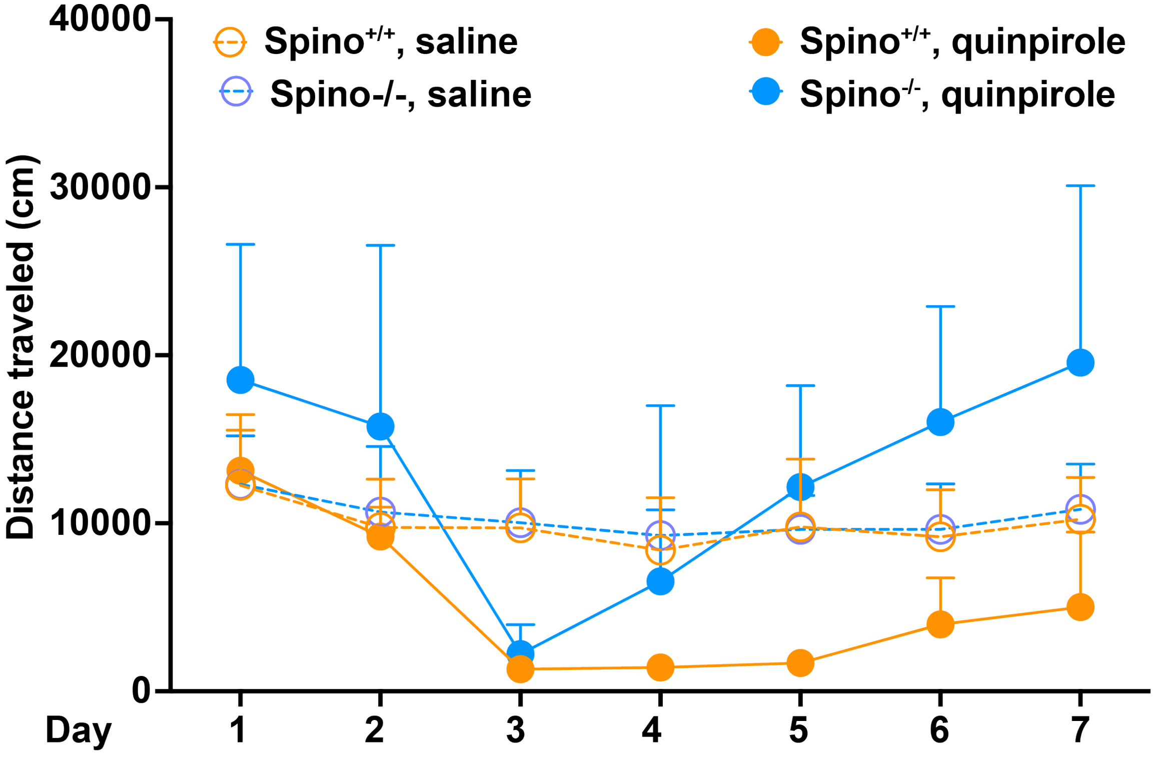

Spino+/+, saline
Spino-/-, saline
Spino+/+, quinpirole
Spino-/-, quinpirole
Distance traveled (cm)
40000
30000
20000
10000
0
Day
1
2
3
4
5
6
7

Figure S5

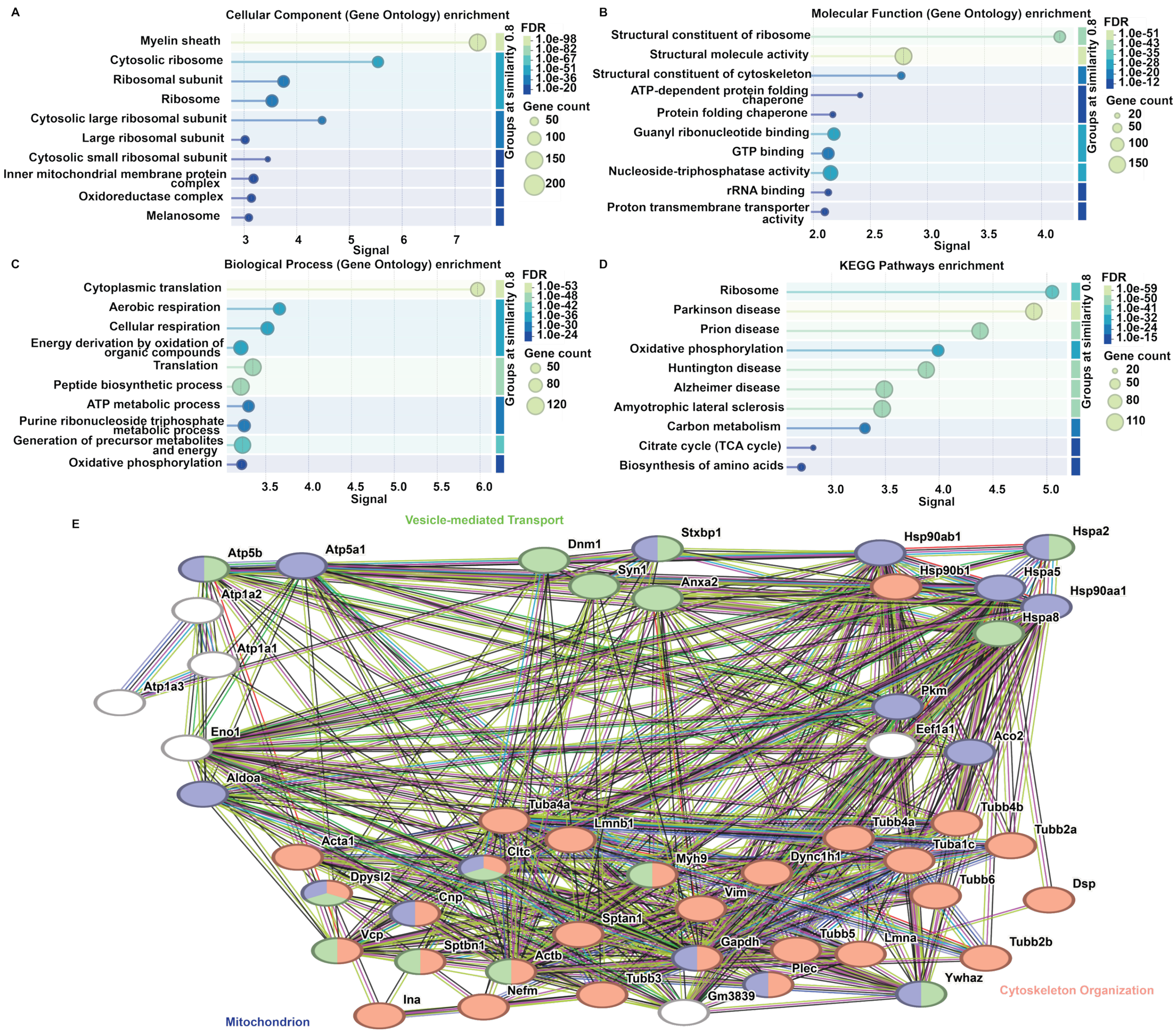

A
Cellular Component (Gene Ontology) enrichment
Myelin sheath
Cytosolic ribosome
Ribosomal subunit
Ribosome
Cytosolic large ribosomal subunit
Large ribosomal subunit
Cytosolic small ribosomal subunit
Inner mitochondrial membrane protein complex
Oxidoreductase complex
Melanosome
3 4 5 6 7
Signal
Groups at similarity 0.8
FDR
1.0e-98
1.0e-82
1.0e-67
1.0e-51
1.0e-36
1.0e-20
Gene count
50
100
150
200
B
Molecular Function (Gene Ontology) enrichment
Structural constituent of ribosome
Structural molecule activity
Structural constituent of cytoskeleton
ATP-dependent protein folding chaperone
Protein folding chaperone
Guanyl ribonucleotide binding
GTP binding
Nucleoside-triphosphatase activity
rRNA binding
Proton transmembrane transporter activity
2.0 2.5 3.0 3.5 4.0
Signal
Groups at similarity 0.8
FDR
1.0e-51
1.0e-43
1.0e-35
1.0e-28
1.0e-20
1.0e-12
Gene count
20
50
100
150
C
Biological Process (Gene Ontology) enrichment
Cytoplasmic translation
Aerobic respiration
Cellular respiration
Energy derivation by oxidation of organic compounds
Translation
Peptide biosynthetic process
ATP metabolic process
Purine ribonucleoside triphosphate metabolic process
Generation of precursor metabolites and energy
Oxidative phosphorylation
3.5 4.0 4.5 5.0 5.5 6.0
Signal
Groups at similarity 0.8
FDR
1.0e-53
1.0e-48
1.0e-42
1.0e-36
1.0e-30
1.0e-24
Gene count
50
80
120
D
KEGG Pathways enrichment
Ribosome
Parkinson disease
Prion disease
Oxidative phosphorylation
Huntington disease
Alzheimer disease
Amyotrophic lateral sclerosis
Carbon metabolism
Citrate cycle (TCA cycle)
Biosynthesis of amino acids
3.0 3.5 4.0 4.5 5.0
Signal
Groups at similarity 0.8
FDR
1.0e-59
1.0e-50
1.0e-41
1.0e-32
1.0e-24
1.0e-15
Gene count
20
50
80
110
E
Vesicle-mediated Transport
Atp5b
Atp5a1
Dnm1
Stxbp1
Hsp90ab1
Hspa2
Atp1a2
Syn1
Anxa2
Hsp90b1
Hspa5
Hsp90aa1
Hspa8
Atp1a1
Atp1a3
Pkm
Eno1
Eef1a1
Aco2
Aldoa
Tuba4a
Lmnb1
Tubb4a
Tubb4b
Tubb2a
Acta1
Cltc
Myh9
Dync1h1
Tuba1c
Dpysl2
Tubb6
Dsp
Cnp
Vim
Sptan1
Tubb5
Lmna
Vcp
Sptbn1
Gapdh
Tubb2b
Actb
Plec
Tubb3
Ywhaz
Ina
Nefm
Gm3839
Cytoskeleton Organization
Mitochondrion

Figure S6

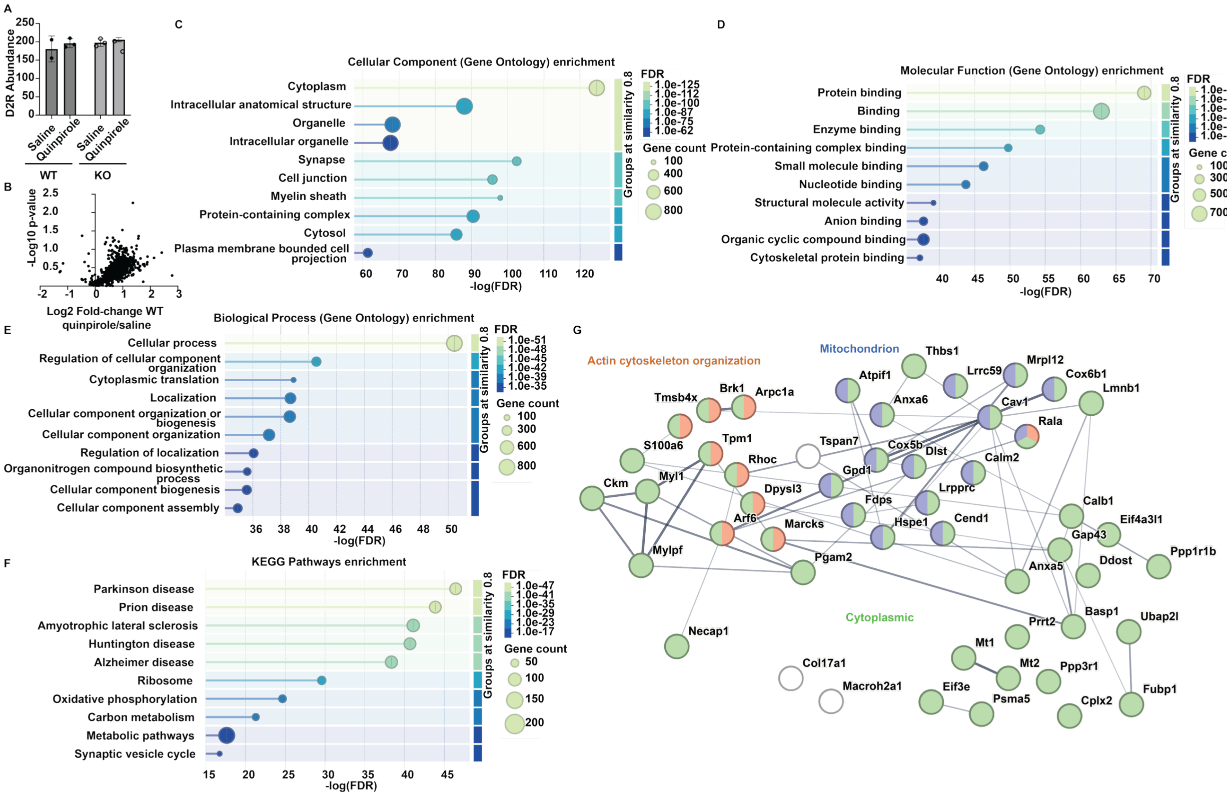
A
D2R Abundance
Saline
Quinpirole
WT
KO
B
-Log10 p-value
Log2 Fold-change WT quinpirole/saline
C
Cellular Component (Gene Ontology) enrichment
Cytoplasm
Intracellular anatomical structure
Organelle
Intracellular organelle
Synapse
Cell junction
Myelin sheath
Protein-containing complex
Cytosol
Plasma membrane bounded cell projection
-log(FDR)
FDR
Gene count
Groups at similarity 0.8
D
Molecular Function (Gene Ontology) enrichment
Protein binding
Binding
Enzyme binding
Protein-containing complex binding
Small molecule binding
Nucleotide binding
Structural molecule activity
Anion binding
Organic cyclic compound binding
Cytoskeletal protein binding
E
Biological Process (Gene Ontology) enrichment
Cellular process
Regulation of cellular component organization
Cytoplasmic translation
Localization
Cellular component organization or biogenesis
Cellular component organization
Regulation of localization
Organonitrogen compound biosynthetic process
Cellular component biogenesis
Cellular component assembly
F
KEGG Pathways enrichment
Parkinson disease
Prion disease
Amyotrophic lateral sclerosis
Huntington disease
Alzheimer disease
Ribosome
Oxidative phosphorylation
Carbon metabolism
Metabolic pathways
Synaptic vesicle cycle
G
Actin cytoskeleton organization
Mitochondrion
Cytoplasmic

Figure S7

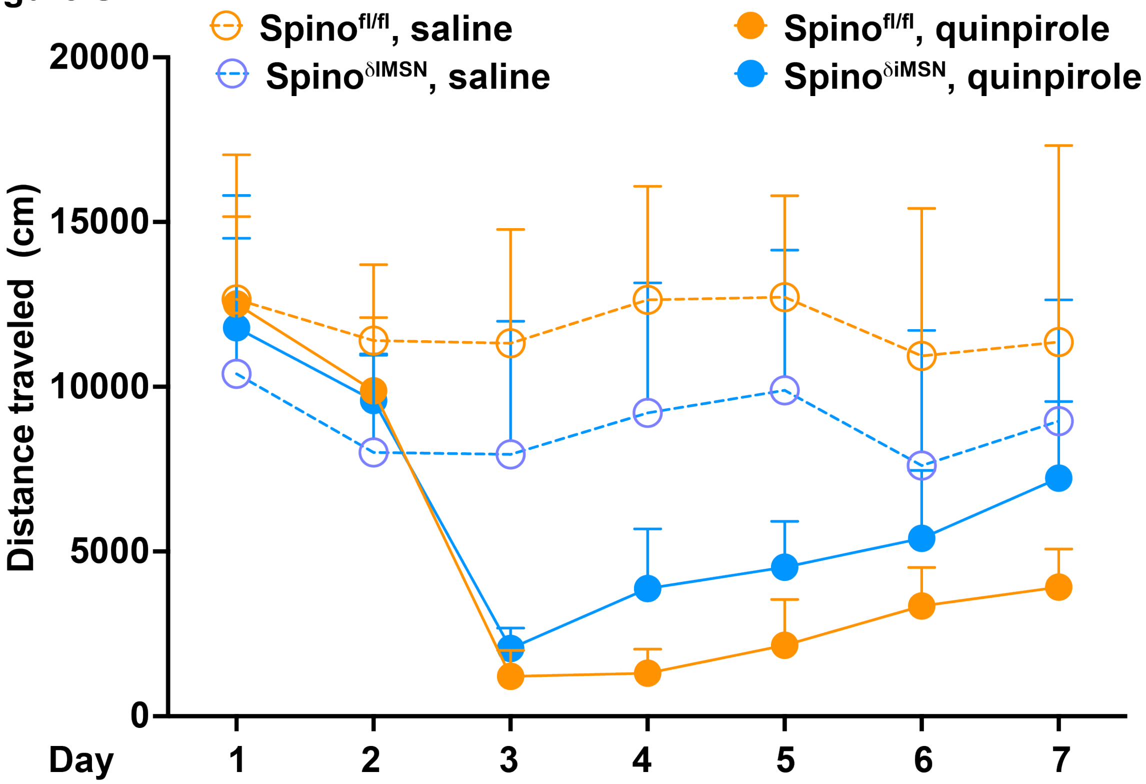

Spino$^{fl/fl}$, saline
Spino$^{\delta IMSN}$, saline
Spino$^{fl/fl}$, quinpirole
Spino$^{\delta iMSN}$, quinpirole
20000
15000
10000
5000
0
Distance traveled (cm)
Day
1
2
3
4
5
6
7

**Figure S8**

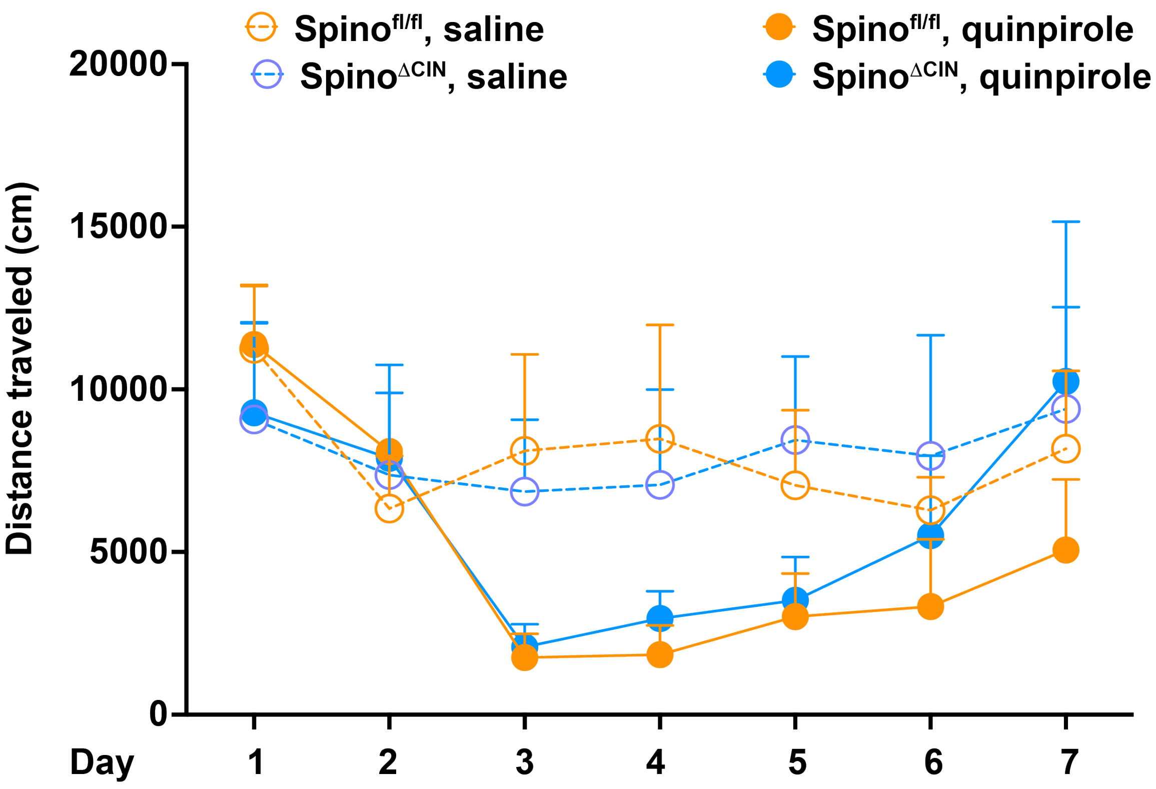

Spino$^{fl/fl}$, saline
Spino$^{\Delta CIN}$, saline
Spino$^{fl/fl}$, quinpirole
Spino$^{\Delta CIN}$, quinpirole
20000
15000
10000
5000
0
Distance traveled (cm)
Day
1
2
3
4
5
6
7

Figure S9

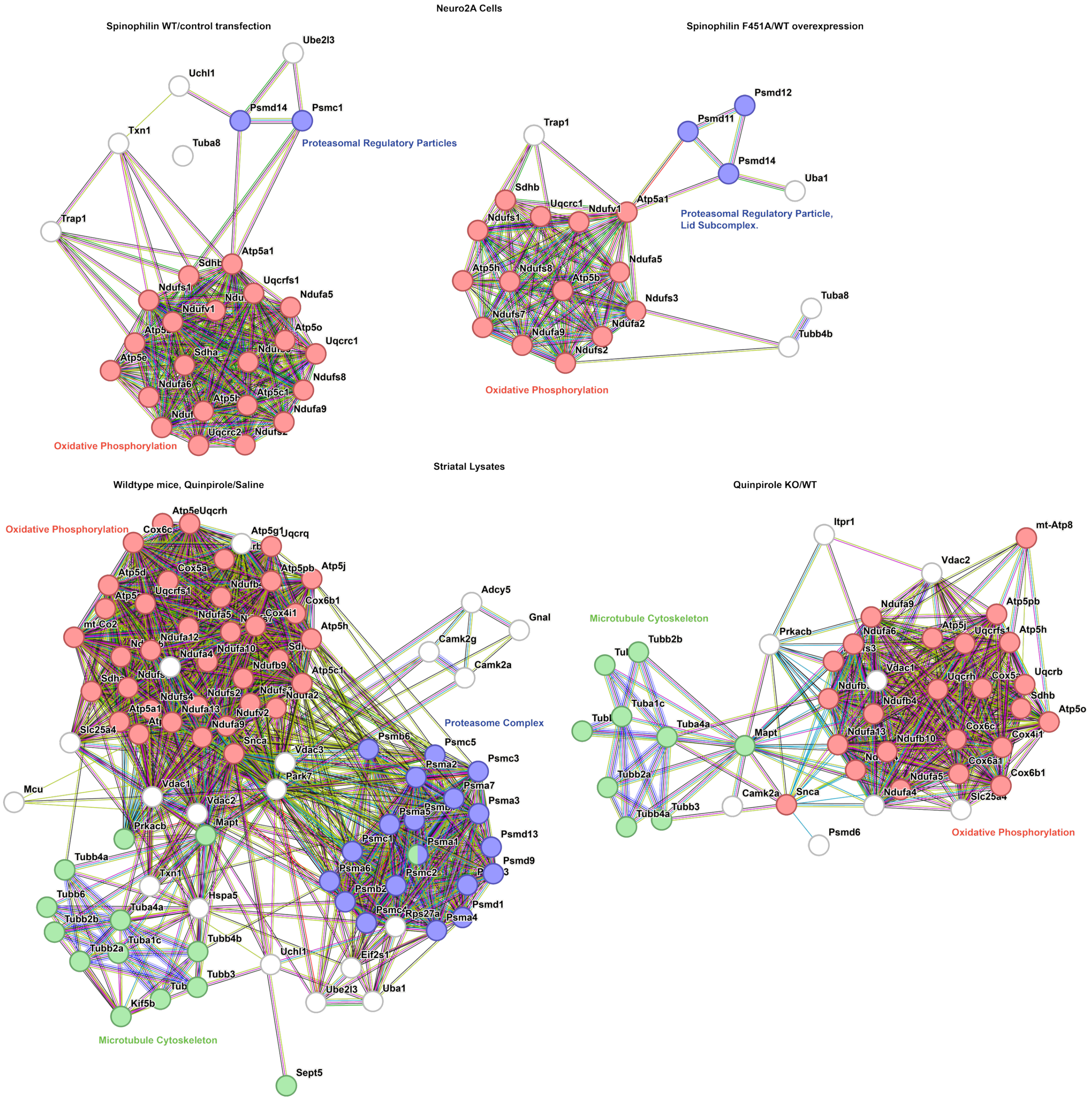
Neuro2A Cells
Spinophilin WT/control transfection
Spinophilin F451A/WT overexpression
Proteasomal Regulatory Particles
Oxidative Phosphorylation
Proteasomal Regulatory Particle,
Lid Subcomplex.
Oxidative Phosphorylation
Ube2l3
Uchl1
Psmd14
Psmc1
Txn1
Tuba8
Trap1
Sdhb
Atp5a1
Ndufs1
Uqcrfs1
Ndufv1
Ndufa5
Atp5o
Uqcrc1
Atp5e
Sdha
Ndufs8
Ndufa6
Atp5b
Atp5c1
Ndufa9
Uqcrc2
Psmd12
Psmd11
Uba1
Uqcrc1
Ndufv1
Atp5h
Ndufs7
Ndufa2
Ndufs3
Ndufs2
Tubb4b
Striatal Lysates
Wildtype mice, Quinpirole/Saline
Quinpirole KO/WT
Oxidative Phosphorylation
Proteasome Complex
Microtubule Cytoskeleton
Microtubule Cytoskeleton
Oxidative Phosphorylation
Atp5e
Uqcrh
Cox6c
Atp5g1
Uqcrq
Atp5pb
Atp5j
Atp5d
Cox5a
Uqcrfs1
Cox4i1
Cox6b1
mt-Co2
Ndufa12
Ndufa4
Ndufa10
Ndufb9
Atp5c1
Ndufs4
Ndufa13
Ndufs2
Ndufa2
Ndufv2
Ndufa9
Slc25a4
Atp5a1
Snca
Vdac3
Park7
Adcy5
Gnal
Camk2g
Camk2a
Psmb6
Psmc5
Psmc3
Psma2
Psma7
Psma3
Psma5
Psmc1
Psma1
Psmd13
Psmd9
Psma6
Psmc2
Psmb2
Psmd1
Rps27a
Psma4
Eif2s1
Uchl1
Ube2l3
Uba1
Sept5
Mcu
Vdac1
Vdac2
Prkacb
Mapt
Tubb4a
Txn1
Hspa5
Tubb6
Tuba4a
Tubb2b
Tuba1c
Tubb2a
Tubb4b
Tubb3
Kif5b
Itpr1
mt-Atp8
Vdac2
Ndufa9
Atp5pb
Ndufa6
Atp5j
Uqcrfs1
Atp5h
Prkacb
Tubb2b
Vdac1
Uqcrh
Uqcrb
Sdhb
Tuba1c
Ndufb4
Atp5o
Mapt
Tuba4a
Ndufa13
Ndufb10
Cox6c
Cox4i1
Cox6a1
Cox6b1
Tubb2a
Ndufa5
Camk2a
Snca
Ndufa4
Slc25a4
Tubb4a
Tubb3
Psmd6
