## Supplementary material for "Striatal spinophilin enhances D2R interaction with cytosolic proteins to mediate persistent D2R agonist-induced locomotor suppression": Key Resources

| **KEY RESOURCES TABLE** |  |  |
| --- | --- | --- |
| REAGENT or RESOURCE | SOURCE | IDENTIFIER |
| Antibodies | | |
| PP1α (E-9) Mouse | Santa Cruz | SC-7482, RRID:AB_628177 |
| Rabbit anti-HA (Y-11) Discontinued | Santa Cruz | Sc-805, RRID:AB_631618 |
| Dopamine D2 Receptor/D2R long isoform | Millipore-Sigma | AB1792P, RRID:AB_11212581 |
| Alexa Fluor 790-conjugated AffiniPure Donkey Anti-Rabbit IgG (H+L) | Jackson-Immuno Research | 711-655-152, RRID:AB_2340628 |
| Alexa Fluor 790-conjugated AffiniPure Donkey Anti-Mouse IgG (H+L) | Jackson-Immuno Research | 715-655-150, RRID:AB_2340870 |
| Alexa Fluor 790-conjugated AffiniPure Donkey Anti-Sheep IgG (H+L) | Jackson-Immuno Research | 713-655-147, RRID:AB_2340754 |
| Alexa Fluor 680 Donkey Anti-Rabbit IgG (H+L) | Invitrogen | A10043, RRID:AB_2534018 |
| Alexa Fluor 680 Donkey Anti-Mouse IgG (H+L) | Invitrogen | A10038, RRID:AB_11180593 |
| Bacterial and virus strains | | |
| N/A |  |  |
| Biological samples | | |
| N/A |  |  |
| Chemicals, peptides, and recombinant proteins | | |
| N/A |  |  |
| Critical commercial assays | | |
| Tandem Mass Tag Pro | ThermoFisher Scientific | A44522 |
| Deposited data | | |
| Raw proteomics data | ProteomeXchange partner MassIVE  ProteomeXchange accession | MSV000098874  PXD067504. |
| Experimental models: Cell lines | | |
| Neuro2A cells | ATCC | RRID:CVCL_0470 |
| Experimental models: Organisms/strains | | |
| C57Bl/6J mice | Jackson Laboratories | RRID:IMSR_JAX:000664 |
| B6.FVB(Cg)-Tg(Adora2a-cre)KG139Gsat/Mmucd | MMRRC | RRID:MMRRC_036158-UCD |
| B6.129S-*Chat^tm1(cre)Lowl^*/MwarJ | Jackson Laboratories, Rossi et al.,^70^ | RRID:IMSR_JAX:031661 |
| Spino^ΔiMSN^ | Morris et al.,^30^ | N/A |
| Spino^ΔCIN^ | This paper | N/A |
| Oligonucleotides | | |
| DRD2 Forward primer - CTCTCCACAGGTGTCCAGGCGGCCGCCATGGTG ATGAAGACGATCATCGCCC | This Paper | N/A |
| GCAACTAGAAGGCACAGTTATTA TAACTTCGTATAGCATACATTATACGAAGTTAT ACAGTGGAGAATCTTCAGAAATG | This Paper | N/A |
| Recombinant DNA | | |
| HA-Spinophilin | Hiday et al.,^17^ | RRID:Addgene 87122 |
| pSF3-ultraID | Kubitz et al.,^24^ | RRID:Addgene  172878 |
| pBS185-iCre | Sauer and Henderson^71^ | RRID:Addgene_11916 |
| pDonr221 | Invitrogen | RRID:Addgene_101410 |
| HA-Spinophilin F451A | This paper | N/A |
| DRD2-TANGO | Kroeze et al.,^68^ | RRID:Addgene_66269 |
| Spinophilin-flox-ALFA-UltraID | Claeboe et al.,^23^ | N/A |
| Software and algorithms | | |
| N/A |  |  |
| Other | | |
| Alexa Fluor 790-conjugated Streptavidin | Jackson-Immuno Research | 016-650-084 |
| Trypsin-resistant streptavidin magnetic beads | ReSyn Biosciences | MR-STP002 |
| Neutravidin Beads | ThermoFisher Scientific | 29200, 29201 |
| Protein G Dynabeads | ThermoFisher Scientific | 10009D |
